# A Cellular Taxonomy of the Adult Human Spinal Cord

**DOI:** 10.1101/2022.03.25.485808

**Authors:** Archana Yadav, Kaya J.E. Matson, Li Li, Isabelle Hua, Joana Petrescu, Kristy Kang, Mor R. Alkaslasi, Dylan I. Lee, Saadia Hasan, Ahmad Galuta, Annemarie Dedek, Sara Ameri, Jessica Parnell, Mohammad M. Alshardan, Feras Abbas Qumqumji, Saud M. Alhamad, Alick Pingbei Wang, Gaetan Poulen, Nicolas Lonjon, Florence Vachiery-Lahaye, Pallavi Gaur, Mike A. Nalls, Yue A. Qi, Michael E. Ward, Michael E. Hildebrand, Pierre-Francois Mery, Emmanuel Bourinet, Luc Bauchet, Eve C. Tsai, Hemali Phatnani, Claire E. Le Pichon, Vilas Menon, Ariel J. Levine

## Abstract

The mammalian spinal cord functions as a community of glial and neuronal cell types to accomplish sensory processing, autonomic control, and movement; conversely, the dysfunction of these cell types following spinal cord injury or disease states can lead to chronic pain, paralysis, and death. While we have made great strides in understanding spinal cellular diversity in animal models, it is crucial to characterize human biology directly to uncover specialized features of basic function and to illuminate human pathology. Here, we present a cellular taxonomy of the adult human spinal cord using single nucleus RNA-sequencing with spatial transcriptomics and antibody validation. We observed 29 glial clusters, including rare cell types such as ependymal cells, and 35 neuronal clusters, which we found are organized principally by anatomical location. To demonstrate the potential of this resource for understanding human disease, we analyzed the transcriptome of spinal motoneurons that are prone to degeneration in amyotrophic lateral sclerosis (ALS) and other diseases. We found that, compared with all other spinal neurons, human motoneurons are defined by genes related to cell size, cytoskeletal structure, and ALS, thereby supporting a model of a specialized motoneuron molecular repertoire that underlies their selective vulnerability to disease. We include a publicly available browsable web resource with this work, in the hope that it will catalyze future discoveries about human spinal cord biology.

## Introduction

The human spinal cord relays, processes, and transforms sensory inputs and descending cues from the brain into sensory, motor, respiratory, and autonomic outputs. These critical processes rely on a diverse array of spinal cord cell types, each with their own functions, molecular repertoires, and vulnerabilities to injury or disease. For example, in hereditary spastic paraplegia, corticospinal, sensory, and spinocerebellar neurons show degeneration (Blackstone, 2018; Bruyn et al., 1994; Schwarz and Liu, 1956); in spinal muscular atrophy spinal motoneurons are primarily affected during development (Arnold and Fischbeck, 2018); and in amyotrophic lateral sclerosis, corticospinal neurons and multiple populations of ventral spinal interneurons die in addition to the signature phenotype involving degeneration of spinal motoneurons (Allodi et al., 2021; Averback and Crocker, 1982; Kawamura et al., 1981; Ravits et al., 2013; Romer et al., 2017; Salamatina et al., 2020; Stephens et al., 2006; Williams et al., 1990). These cell types have been extensively studied in model organisms, including molecular profiling of all spinal cord cell types at the single-cell level in the mouse spinal cord (Häring et al., 2018; Osseward et al., 2021; Rosenberg et al., 2018; Russ et al., 2021; Sathyamurthy et al., 2018; Zeisel et al., 2018). However, technical obstacles and limited access to high quality tissue specimens have prevented the full application of single cell approaches to study human spinal cord biology. Thus, prior work has only been done on limited cell types or in human fetal tissue (Rayon et al., 2021; D. Zhang et al., 2021; Q. Zhang et al., 2021).

To characterize the cell types of the adult human lumbar spinal cord, we used recently optimized tissue extraction methods on spinal cords from organ donor subjects and performed single nucleus RNA-sequencing of over 50,000 nuclei. We identified 64 unique clusters including 29 non-neuronal populations and 35 neuronal populations and validated many of the predicted expression patterns with independent spatial transcriptomics profiling on an independent sample. We established a comprehensive taxonomy of the neuronal clusters, compared them with their mouse counterparts, and created a publicly available browsable interface as a resource for the field (https://vmenon.shinyapps.io/hsc_biorxiv/). Finally, we performed a focused analysis on the transcriptional profile of spinal motoneurons, identifying a molecular signature that could underlie their selective vulnerability in neurodegenerative disease.

## Results

We obtained post-mortem lumbar spinal cord tissue from seven donor transplant cases (Fig. 1a and Data File Table S1), using neuroprotective conditions, such as body chilling and perfusion with a high magnesium solution, rapid collection, and flash freezing of tissue immediately in the operating room (see Methods). Single nuclei were isolated and profiled, resulting in a dataset of 55,420 nuclei that passed quality control filtering; with median detection of 2,187 genes detected per nucleus. Initial clustering of all nuclei clearly distinguished the major known cell classes present in spinal cord tissue, including oligodendrocytes and their precursors and progenitors, meningeal cells, astrocytes, endothelial and pericyte cells, microglia, and neurons; the latter included glutamatergic neurons, GABAergic/glycinergic neurons, and motoneurons. Comparison of these cell classes to our prior work in the mouse spinal cord (Russ et al., 2021) revealed substantial overlap in cellular signatures as well as notable differences. For example, oligodendrocytes accounted for a larger proportion of the nuclei in the human dataset. This observation is consistent with the larger ratio of white matter to gray matter area in human versus mouse spinal cords (Supplemental Fig. S1A) and could reflect the relative expansion of long axon tracts linking the brain and spinal cord in humans. To determine whether the overall proportions of cells classes that we observed in the sequencing dataset reflected *in vivo* tissue composition, we analyzed the prevalence of oligodendrocytes, astrocytes, microglia, and neurons in adult human lumbar spinal cord tissue. We found similar proportions for neurons, astrocytes, microglia, and oligodendrocytes (Fig. 1D, p = 0.67, p = 0.33, p = 0.06, p = 0.06) in tissue versus dissociated nuclei. Overall, the major cell classes in the sequencing dataset showed clear segregation of previously reported markers for these cell types, thus allowing for further investigation within each of these broad classes (Fig. 1B, Supplemental Fig. S2-S6), as described below.

**Fig. 1:**
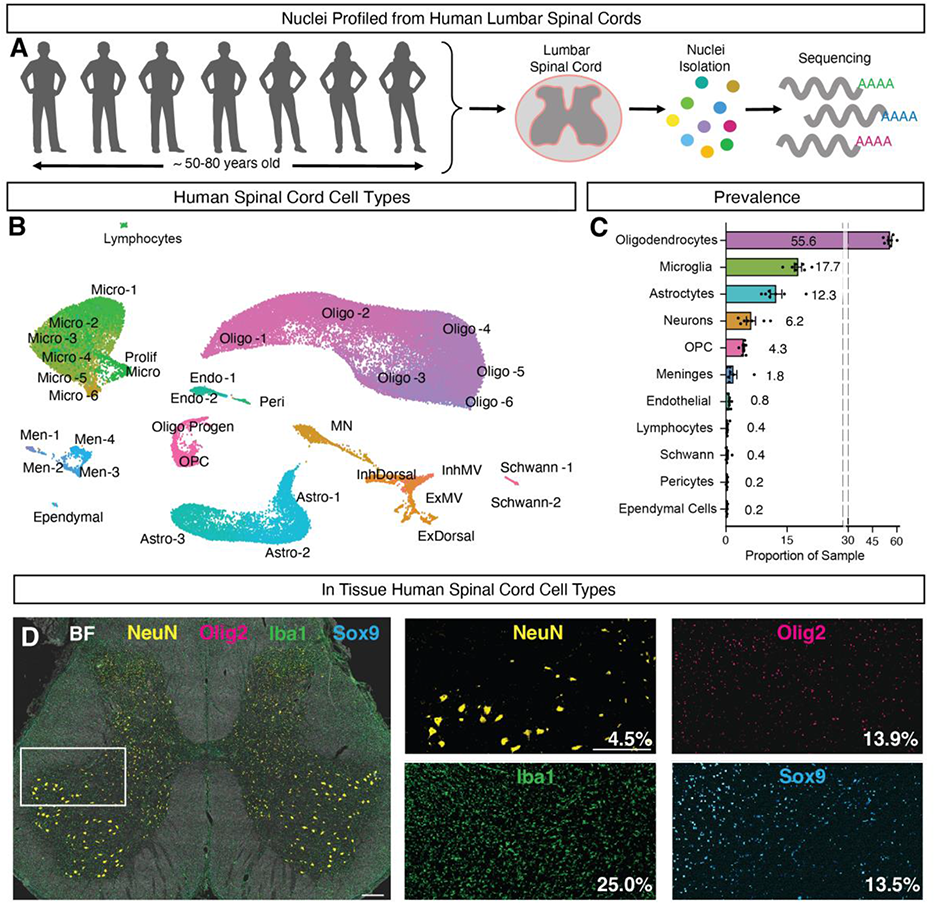
A single cell catalog of the human spinal cord reveals the gene expression signature of human motoneurons. **A,** Lumbar spinal cord tissue was obtained from seven subjects (male and female, ∼50-80 years old) and processed for single nucleus RNA sequencing. **B,** UMAP plot showing the major cell types of the human spinal cord, each in separate color. Cells of the oligodendrocyte lineage are shown in pink/purple and include oligodendrocyte precursor cells (OPC), progenitors (Oligo Progen), six groups of oligodendrocytes (Oligo-1 through Oligo-6), as well as two populations of Schwann cells (Schwann-1 and –2). Microglia cells are shown in green and includes a putatively proliferating population (Prolif Micro) and six groups of microglia (Micro-1 through Micro-6). Astrocytes are shown in turquoise and include three populations (Astro-1 through Astro-3). Meninges are shown in blue and include four populations (Men-1 through Men-4). Vascular cells are shown in teal and include two groups of endothelial cells (Endo-1 and –2) and pericytes (Peri). Ependymal cells are shown in teal. Neurons are shown in orange and include five broad classes based on their neurotransmitter status and putative location: motoneurons (MN), excitatory dorsal neurons (ExDorsal), inhibitory dorsal neurons (InhDorsal), excitatory mid neurons (ExM), excitatory ventral neurons (EV), and inhibitory mid neurons (InhM) and inhibitory ventral neurons (InhV). **C,** Bar plot showing the proportion of a given cluster in each donor (N=7). Error bars are ± s.e.m. **D,** Multiplex immunohistochemistry of the lumbar human spinal cord, stained for NeuN (yellow), IBA1 (green), SOX9 (turquoise), OLIG2 (pink). Brightfield (BF) is shown in white. Percent of DAPI+ cells expressing NeuN, OLIG2, IBA1 and SOX9 are noted in the bottom right corner of each inset (N = 2). Scale bars are 500 μm. Accompanying bar plots are in Supplemental Fig. S7.

### Glial and Support Cell Populations of the Adult Human Lumbar Spinal Cord

Clustering of non-neuronal classes identified specific subpopulations that we identified by homology with related mouse cell types (Russ et al., 2021) and that we partially validated using spatial transcriptomics on post-mortem tissue from an independent donor (Fig. 2). Amongst oligodendrocytes and related populations, we observed two populations of Schwann cells that were detected at the edges of the spinal tissue in the dorsal root entry zone, a population of oligodendrocyte precursor cells and related progenitors, as well as six populations of oligodendrocytes that were distributed over the entire spinal cord tissue with a bias for the white matter tissue, as expected (Fig. 2A-C, Supplemental Fig. S6). Amongst microglia, we observed six populations, including a putative proliferative type characterized by expression of POLQ, TOP2A, and MKI67 (Fig. 2D-F, Fig. 2M, Supplemental Fig. S6). In prior work on adult mouse spinal cord cell types, proliferative microglia were not observed in the healthy spinal cord, including in mature adult (5-6 month old) animals (Matson et al., 2021; Squair et al., 2021). We therefore analyzed post-mortem tissue from three independent organ donor subjects not included in the single nucleus RNA-sequencing dataset to confirm the existence of this population existed in intact tissue. Indeed, we found that 23 percent of microglia in tissue co-expressed the proliferative marker Ki67 (Supplemental Fig. 7, 25% of cells were IBA1+, with 5.77% of cells double positive for IBA1 and Ki67). Whether this reflects normal human biology, is an aging-induced phenotype, or due to peri-mortem changes remains to be determined. Amongst astrocytes, we identified three populations, including one that localized to the white matter in the spatial transcriptomics data, and two that were localized to the gray matter. These gray matter astrocytes populations (ASTRO-2 and ASTRO-3) were enriched for genes involved in neural metabolism and signaling including the GABA transporter SLC6A11, the AMPA receptor regulator SHISA9, and the synaptic adhesion protein TENM2 (Fig. 2G-I, Fig. 2M, Supplemental Fig. S6). By contrast, the white matter astrocyte population was enriched for CD44, CPAMD8 and AQP4. Finally, amongst support cells, we identified two endothelial cell populations, one pericyte population, four populations of meningeal cells, a group ependymal cells and a group of lymphocytes (Fig. 2J-L).

**Fig. 2:**
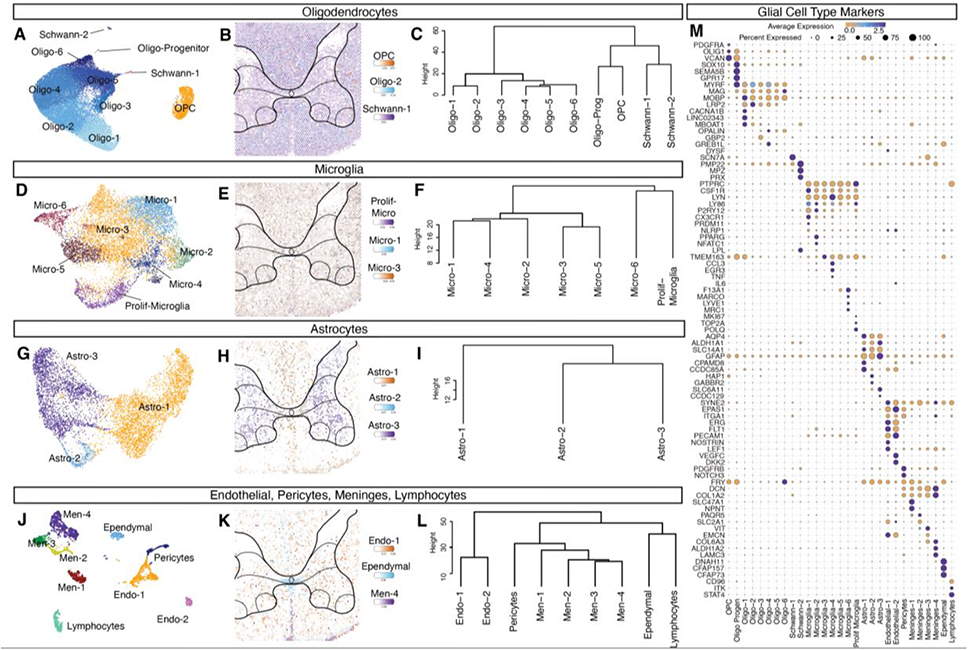
Glial and support cell types in the human spinal cord. Glial cell types including **A-C,** Oligodendrocytes, **D-F,** Microglia **G-I,** Astrocytes **J-L,** Endothelial cells, ependymal cells, pericytes and lymphocytes. For each cell type, a UMAP shows the subtypes, a spatial feature plot shows Cell2Location data, and a dendogram depicts the relationships between the subtypes. Individual spatial transcriptomic spatial feature Cell2Location plots can be found in Supplemental Figure S6. Dendograms were calculated using the top 2,000 highly variable genes from each population (Ward’s method). **M,** Dot plot of markers for glial subtypes showing average expression and percent expressed. Average expression ranges from low (orange) to high (purple).

### Neuronal Atlas of the Adult Human Lumbar Spinal Cord

To characterize the neuronal populations of the adult human lumbar spinal cord, we sub clustered the neuronal nuclei and identified 35 groups. These include a large population of spinal motoneurons (described in greater detail below) and 33 glutamatergic (defined by the expression of SLC17A6) or GABA/glycinergic populations (defined by expression of GAD1, GAD2, PAX2, and SLC6A5). Importantly, each of these populations contained nuclei from each of the seven donors (Fig. 3A, Supplemental Fig. S8A). (We also observed one neuronal cluster that was defined by expression of immediate <u>e</u>arly response <u>g</u>enes (IEG), though it is unclear whether this reflects neuronal activity/stress during the patient’s life or post-mortem artifacts.) Interestingly, amongst the most differentially enriched genes between putative excitatory and inhibitory cell types, we observed a pair of calcium channel regulatory subunits (CACNA2D1 and CACNA2D3) and a pair of self-avoidance adhesion molecules (DSCAM and DSCAML1), both of which are conserved in mice (Russ et al., 2021). This latter signature raises the possibility that excitatory-inhibitory network balance may be achieved partly through self-avoidance control of synaptic connectivity.

**Fig. 3:**
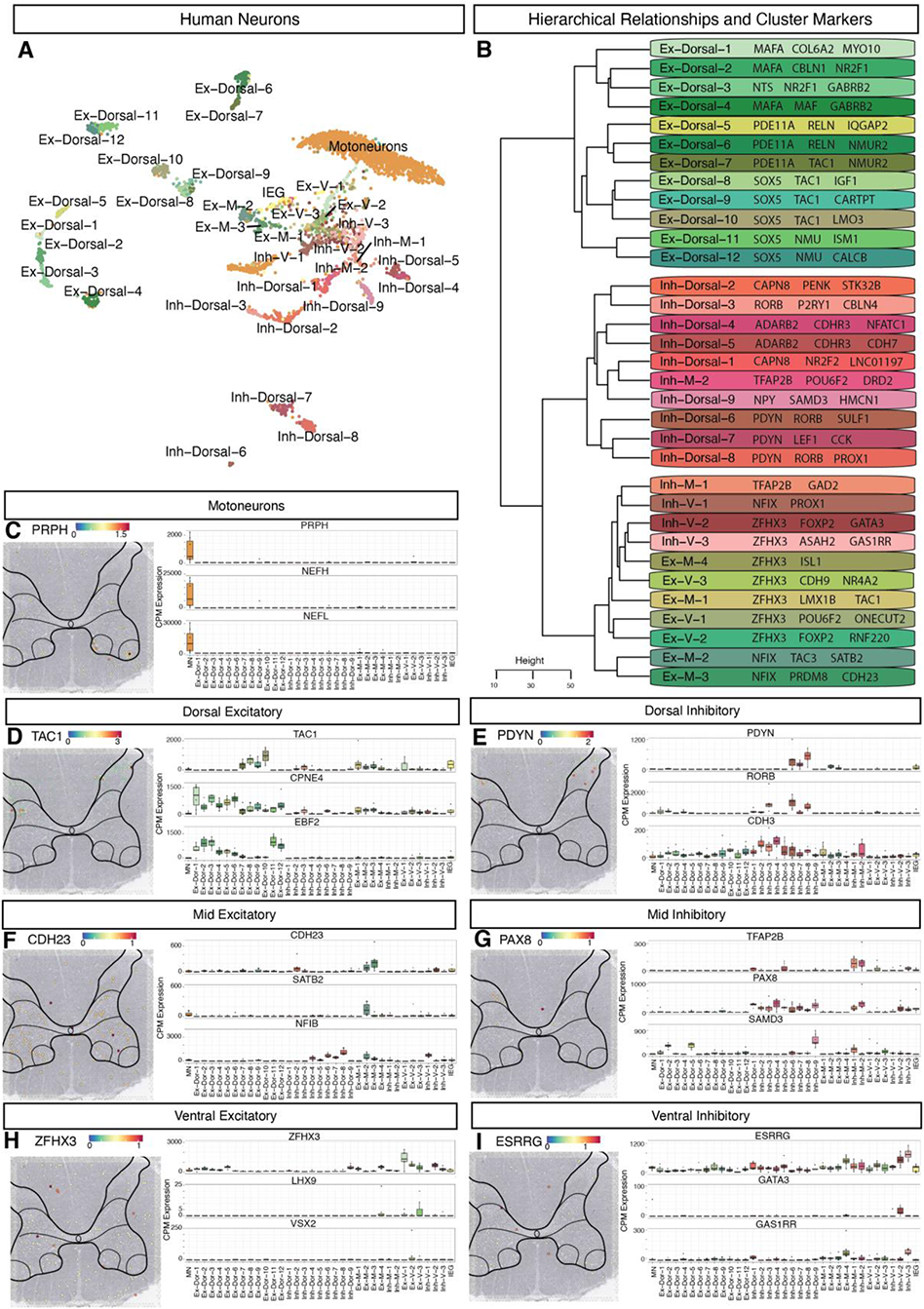
Neuronal cell types in the human spinal cord. **A,** UMAP plot of human spinal neurons showing 35 refined populations. **B,** Dendogram showing relationship of neuronal subtypes, calculated using the top 2,000 highly variable genes (Ward’s method). For each cluster, 2-3 top genes are shown. **C-I,** For each class of spinal cord neuron (Motoneurons, Dorsal Excitatory, Dorsal Inhibitory, Mid Excitatory, Mid Inhibitory, Ventral Excitatory and Ventral Inhibitory) spatial feature plots shows the expression of a marker in tissue and box plot shows per-cluster and per-sample expression (Counts per Million) of 3 marker genes. Box plots show average expression from each donor (N = 7). Outliers are plotted with a dot. **D,** Dorsal excitatory markers include TAC1, CPNE4, and EBF2. **E,** Dorsal inhibitory markers include PDYN, RORB, and CDH3. **F,** Mid excitatory markers include CDH23, SATB2, and NFIB. **G,** Mid inhibitory markers include TFAP2B, PAX8, and SAMD3. **H,** Ventral excitatory markers include ZFHX3, LHX9, and VSX2. **I,** Ventral inhibitory markers include ESRRG, GATA3, and GAS1RR. Spatial transcriptomic gene expression is colored from purple (low) to red (high).

Given that the function of spinal cord neurons is highly related to their anatomical location, we explored the spatial distribution of the 33 excitatory and inhibitory populations. We used a combination of comparison to spatial transcriptomics data for key marker genes and comparison with data from macaque and mouse to assign putative locations for each population, sorting them into general categories of dorsal, mid, and ventral cell types (Fig 3B). We then adopted a nomenclature for these cell types that references both their putative location and neurotransmitter status. To reveal the overall organization of human lumbar spinal neuronal populations, we analyzed their relationships with three different approaches: correlation of their gene expression profiles (Fig 5C), proximity in gene expression-derived principal component space (Supplemental Fig. S12), and their separability by silhouette scoring (silhouette coefficient; Fig. 4, Supplemental Fig. S9B) and random-forest based machine learning classifier (Supplemental Fig. S10). Each of these methods revealed the same patterns: (1) location in either the dorsal or mid/ventral domain was the primary factor in overall cell type organization, (2) putative dorsal neuron populations were well separated from each other into robust, distinct clusters with highly significant differential molecular markers, and (3) mid and ventral neuronal clusters were in less clearly distinct with partially overlapping gene expression profiles, were closer in principal component space and had lower accuracy in post-hoc classification. These findings are similar to trends observed in mouse spinal neurons, establishing dorsal-ventral location as the conserved, core organizational axis of spinal neuron variability in both species.

**Fig. 4:**
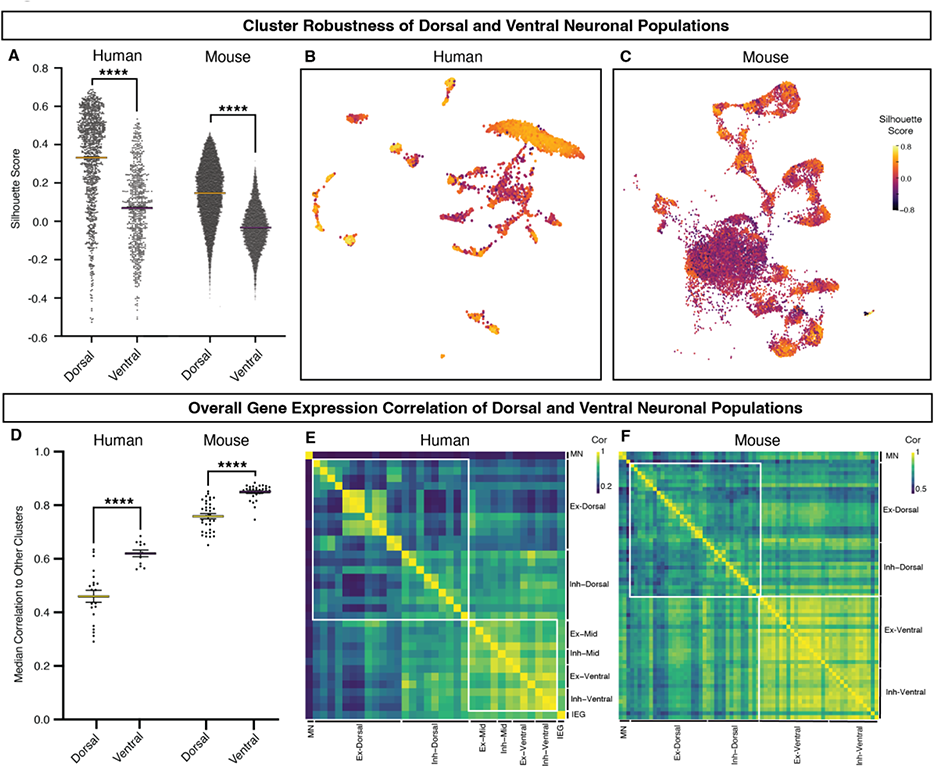
Overall organization of human and mouse lumbar spinal neuronal populations. **A,** The relationship between dorsal and ventral neurons in the human and mouse spinal cord neurons compared using a silhouette score, with values ranging from –1 to 1 (where a high value indicates that clusters are significantly distinguished from one another). (Individual cluster silhouette scores are shown in Supplemental Fig. S8B.) Two-way ANOVA and Mann Whitney test for human and mouse dorsal vs ventral distributions are as follows. P < 0.0001, ****. B-C, UMAP of human neurons (**B**) and mouse neurons (**C**) colored by Silhouette score— purple (low) to yellow (high). **D,** Median correlation of a cluster to other clusters, as calculated by Pearson’s Correlation using the top 2,000 highly variable genes. Two-way ANOVA and Mann Whitney test for human and mouse dorsal vs ventral distributions are as follows. P < 0.0001, ****. **E-F,** Heatmap correlation plot of the human spinal cord neurons (**E**) and mouse spinal cord neurons (**F,** Russ et al. 2021). Correlation is colored from purple (low) to yellow (high).

**Fig. 5:**
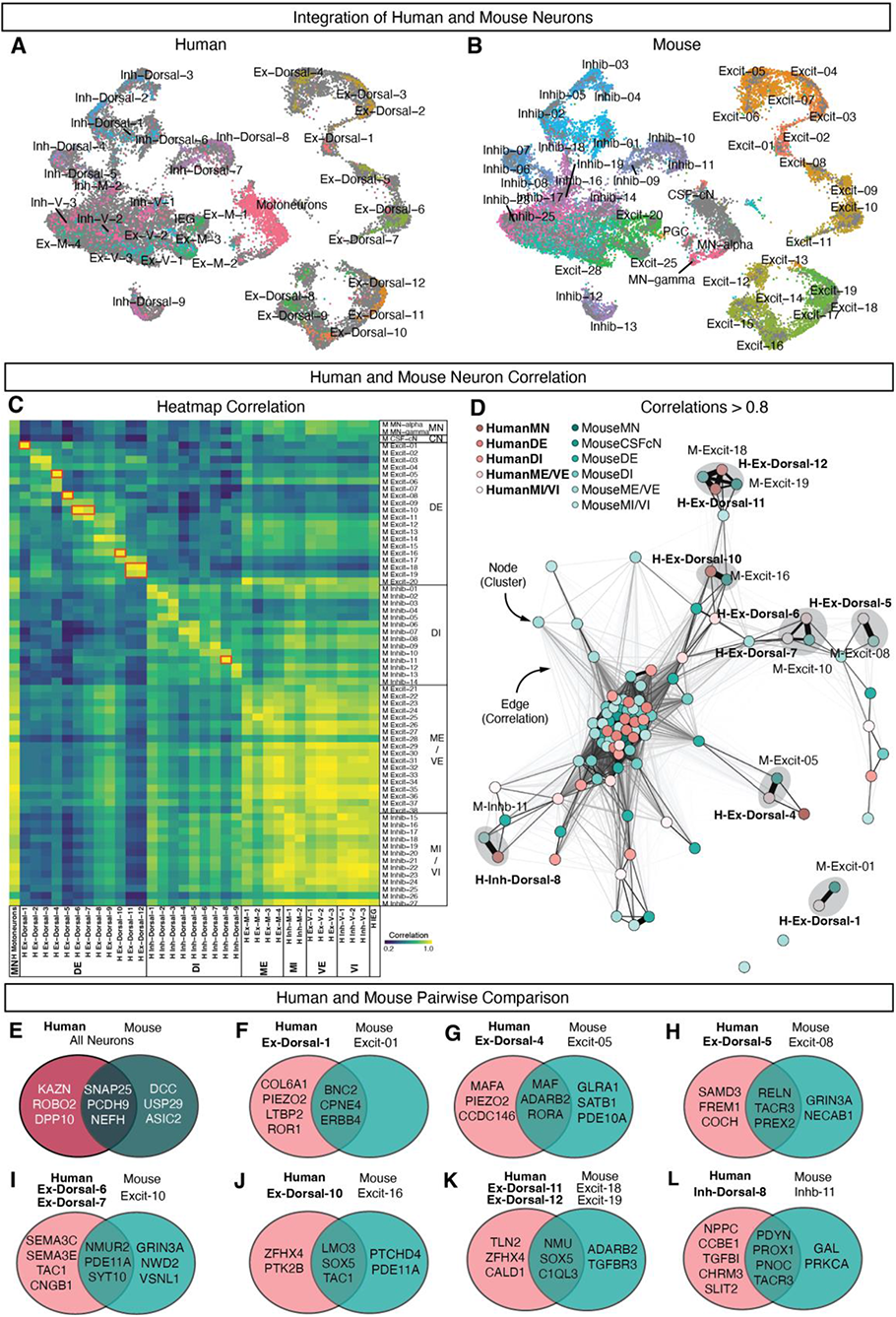
Integration of human and mouse spinal cord neurons. **A,** UMAP plot of human spinal neurons. **B,** UMAP plot of mouse spinal neurons (Russ et al. 2021). **C,** Heatmap correlation plot of the human spinal cord neurons compared to mouse populations (Russ et al. 2021). Correlation is colored from purple to yellow and was calculated using the top 2,000 highly variable genes. Red boxes highlight 7 pairs of clusters shown in **E-L**. Human clusters are bolded and mouse clusters are in regular font. **D,** A forced graph (quotient) showing neuronal clusters as nodes connected by edges. Edges represent correlations greater than 0.8 between human and mouse neuronal clusters. Line thickness and distance indicates correlation value, with greater correlations having a thicker and shorter line. Human neuronal clusters are bolded and shown in shades of pink. Mouse neuronal clusters are shown in shades of blue. Grey circles highlight 7 pairs of clusters shown in **E-L. E-L,** Venn-diagrams represent differentially expressed genes between human and mouse pairs, as well as genes shared by a pair of clusters vs all other neurons. Top genes enriched in the human neurons from each pair are shown in the pink circle, and top genes enriched in mouse neurons in the blue circle. Genes enriched in the human and mouse pair compared to all other neurons are shown in the intersection of the venn-diagram.

We next performed a detailed comparison of individual human and mouse spinal cord neuronal populations by integrating our work with prior harmonized datasets from postnatal mouse tissue (Fig. 5A-B). We found that, overall, human neurons were enriched for KAZN, ROBO2 and DPP10 while mouse neurons were enriched for DCC, USP29, and ASIC2 (Fig. 5E). There was good correspondence between the two datasets, with pairs of human-mouse dorsal clusters showing high correlations and specific relationships, while ventral clusters showed broader overall similarity (Fig 5C). We used a network perspective on cluster relatedness to highlight human and mouse cell types pairs with particularly high conservation and analyzed these further (Fig. 5D). As examples: (1) Human Ex-Dorsal-4 is highly homologous to mouse Excit-05, a member of the Maf family located in lamina III-IV which is associated with corrective reflexes and light touch processing (Fig. 5G). Both the human and mouse clusters are enriched for MAF, ADARB2, and RORA, while the human cluster is also enriched for MAFA (found in the spatial transcriptomics data in the deeper region of the dorsal horn) and the mechanosensitive protein PIEZO2, which may confer evolutionarily novel functions on this population. (2) Human Inh-Dorsal-8 was highly homologous to mouse Inhib-11, a member of the Pdyn family located in lamina I-III and associated with mechanical allodynia pain symptoms (Fig. 5L). Both clusters were enriched for the neuropeptides PDYN and PNOC (both found in the spatial transcriptomics data over the dorsal horn), as well as PROX1 and TACR3. The human cluster was enriched for the neuropeptide NPPC while the mouse cluster was enriched for the neuropeptide Gal. In the future, such cross-species cell type relationships can be used to propose behavioral functions for a broad range of human neuronal populations.

### Human motoneurons are defined by genes related to cell structure, cell size, and ALS

We next sought to use this cellular and molecular resource to study the gene expression profile of human motoneurons and to determine whether their molecular repertoire provided insight into their selective vulnerability in diseases such as ALS and SMA.^1^ We examined the top 50 marker genes that distinguished the motoneuron cluster from other human spinal neurons. To determine whether these genes were enriched in motoneurons in spinal cord tissue, we assessed the distribution of the entire predicted gene signature using the spatial transcriptomics dataset from an independent donor subject. Indeed, this signature was strongly enriched in the most ventral spinal tissue, confirming the overall pattern of motoneuron marker genes (Supplemental Fig. S13). Overall, the motoneuron markers included those involved in acetylcholine synthesis and function (SLC5A7 and ACLY), as expected, but surprisingly were dominated by three partially overlapping sets of genes: (1) those involved in cytoskeletal structure, (2) neurofilament genes related to cell size, and (3) those that are directly implicated in ALS pathogenesis (Fig. 6A).

**Fig. 6.**
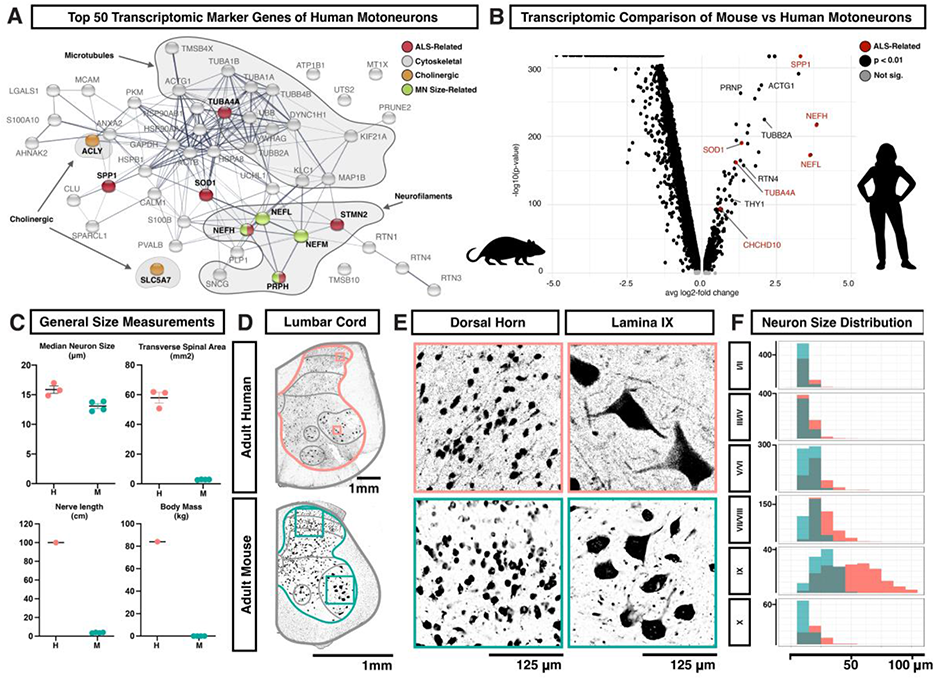
Human motoneurons are characterized by genes associated with ALS, cell structure, and increased cell size. **A,** Association network plot constructed using the String protein database for the top 50 marker genes of human motoneurons. Genes related to cholinergic neurotransmission are shown in orange, genes related to ALS are shown in red, and genes whose over-expression in mice causes enlargement and/or degeneration of motoneurons are shown in green. Families of genes related to the microtubule or neurofilament cytoskeletal components are highlighted by gray. **B,** Volcano plot showing the distribution of genes enriched in either lumbar motoneurons from adult mice or lumbar motoneurons from adult humans, with several significant genes of interest labeled, including genes related to ALS (red). Genes are plotted by the average change in expression (avg log_2_-fold change) and by the statistical strength of the difference (-log_10_(p-value). Insignificant genes are shown in gray and genes that are significantly different are shown in black or red. **C,** Gross anatomical and neuronal measurements of the human (H) and mouse (M) lumbar spinal cords. Measurements include median neuron size (µm), transverse area of the spinal cord (mm^2^), maximum nerve length (cm), and body mass (kg). **D,** Transverse sections of one side of the adult lumbar human (above) and mouse (below) spinal cords, with antibody labeling for NeuN. Images are representative of data from three subjects. Scale bars are 1 mm. Boxes indicate the regions shown in panel E. Gray lines indicate the laminar/regional boundaries used in panel F and were based on prior work (Routal and Pal, 1999; Schoenen, 1991; “The Human Nervous System,” 2004; Watson et al., 2009). **E,** Higher magnification view of NeuN labeled spinal neurons from panel **D** in the human (above) and mouse (below). The left-side images are from the dorsal horn and the right-side images are of putative motoneurons in lamina IX. Scale bars are 125 µm. **F**, Histogram showing the count distribution of neuron Feret distance (maximum caliper, similar to diameter) in human (pink) and mouse (teal) across the different lamina regions of the adult lumbar spinal cord. Measurements are given in µm and the count scale is shown at the right of each plot. Bonferroni-adjust Wilcox test p-values and Bhattacharyya Coefficients (BC) for human vs mouse distributions are as follows. I/II: p=7.5e-27, BC=0.93, III/IV: p=4.0e-12, BC=0.96, V/VI: p=3.2e-30, BC=0.89, VII/VIII: p=5.7e-49, BC=0.80, IX: p=1.6e-19, BC=0.71, X: p=9.5e-10, BC=0.92.

Cytoskeletal components were the most abundant category of motoneuron marker gene and the most enriched gene ontology (GO) terms, including GO annotation clusters related to microtubules (p=0.000009) and axon structure and neurofilaments (p=0.000018) (Data File Table S4). The marker genes that were structural components of neurofilaments (NEFL, NEFM, NEFH, and PRPH) have been directly linked to cell size, axon diameter, and degeneration (Beaulieu et al., 1999; Côté et al., 1993; Gama Sosa et al., 2003; Marszalek et al., 1996; Xu et al., 1993a; 1993b), providing a potential link between human motoneuron gene expression and cellular phenotype. Amongst ALS-related motoneuron marker genes, there were both cytoskeletal genes (NEFH, PRPH, TUBA4A, and STMN2), as well as genes that are not directly linked to cellular structure (SOD1, OPTN, and SPP1).

We further examined the expression of a panel of ALS-related genes compiled from the literature (Brown and Al-Chalabi, 2017; Castellanos-Montiel et al., 2020; Gregory et al., 2020; Klim et al., 2019; Morisaki et al., 2016; Taylor et al., 2016; Theunissen et al., 2021; Yamamoto et al., 2017) across human spinal cord cell types. In addition to the genes above, we found that CHCHD10 and KIF5A were enriched in spinal motoneurons, extending this signature profile (Supplemental Fig. S15 and S17). We also observed enriched expression of SPP1, FUS, and C9ORF72 in microglia and STMN2, and TUBA4A in an excitatory mid-population (Ex-M-1, Supplemental Fig. S16, S17). TARDBP was not detected at sufficient levels in the dataset to characterize its expression pattern.

The enriched expression of neurodegeneration-associated genes in human motoneuron transcriptomics may have been partly due to the age of the study donors. We examined expression of ALS-related genes in a dataset of human embryonic spinal cord cell types (Rayon et al., 2021) and found low levels of gene expression (i.e. NEFH and TUBA4A), moderate but broad cell type expression (i.e. OPTN and PRPH), or high and ubiquitous cell type expression (i.e. SOD1 and STMN2) (Supplemental Fig. S17). Thus, the enrichment of ALS-related genes in human motoneurons was not apparent in newly formed motoneurons but likely emerged at some point during motoneuron maturation or aging. Finally, to test whether this expression profile reflected a non-specific enrichment of degeneration-associated genes in human motoneurons with age, we compared the expression of genes for multiple neurodegenerative diseases, including those with age-related associations, across human spinal cord cell types. This analysis revealed a specific association of ALS-related gene expression in human motoneurons (Supplemental Fig. S21)

To determine whether ALS-related genes are also enriched in motoneurons in mice, the major animal model for studying the genetic basis of neurodegenerative disease, we compared the human data to prior single nucleus sequencing data from lumbar skeletal motoneurons from adult mice (Alkaslasi et al., 2021). We found that prominent ALS-related genes were enriched and were expressed at higher levels specifically in the human motoneurons as compared to mouse motoneurons (Fig. 7C). To determine if this enrichment is unique to motoneurons, we examined the analysis of a recent study on conservation in human brain gene expression patterns (Pembroke et al., 2021) and found that three genes of interest (SOD1, TUBA4A, OPTN) had a significantly higher mean human to mouse divergence score than other assayed genes (mean score of 0.587 ± 0.19 versus 1,426 other genes with mean 0.320 ± 0.123, p=0.0002).

**Fig. 7.**
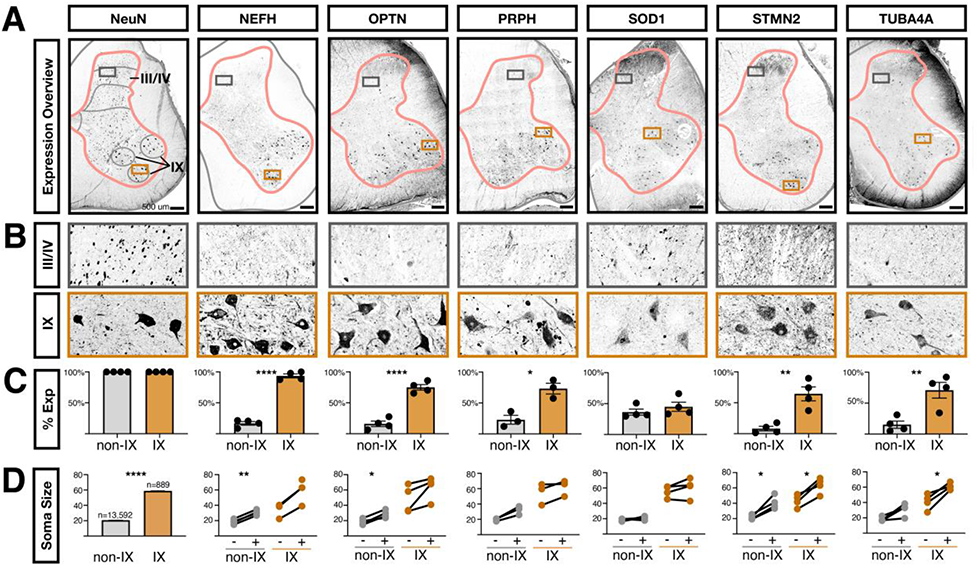
ALS-related proteins are enriched in human motoneurons. **A,** Antibody staining on adult human lumbar spinal cord against NeuN (RBFOX3 gene, general neural marker) and the ALS-related genes NEFH, OPTN, PRPH, SOD1, STMN2, and TUBA4A. Gray matter outlines are shown in pink and boundaries of lamina I/II, III/IV, V/VI, VII/VIII, IX, and X are shown in gray. Boxes indicate the enlarged images in panel. **A,** Images are representative of data from three subjects (two male and one female). Scale bars are 500 µm. **B,** Inset of the images in panel **A,** from the boxed region in laminae III/IV or lamina IX. The width of the insets is 500 µm. **C,** Quantification of the percent of NeuN+ neurons that co-expressed the indicated proteins in either all neurons not in lamina IX (non-IX) or those in lamina IX. The mean ± s.e.m. are shown. The plotted values and number of cells counted in each subject and category are available in Data File Table S5). Paired t-test results are shown where * indicates p < 0.05, ** indicates p < 0.005, **** indicates p < 0.0001. **D,** The sizes of NeuN+ neurons are shown for each indicated protein. For NeuN, 100% of cells were positive, by definition, and the total counts and sizes (mean ± s.e.m.) are shown for neurons not in lamina IX (non-IX) or those in lamina IX. For all other indicated proteins, the Feret distance sizes are shown for all neurons that did not (-) or did (+) express the indicated protein (mean Feret distance in µm). Each line joins values within one subject. There is an unpaired value for NEFH because we did not detect neurons in lamina IX that did not express NEFH. The plotted values and number of cells measured in each subject and category are available in Data File Table S5. Paired two-tailed t-test p-values, after Benjamini-Hochberg FDR correction, are shown where * indicates p < 0.05, ** indicates p < 0.005. **** indicates p < 0.0001.

### Cell Size and Protein Expression in Human Lumbar Motoneurons

Why might human motoneurons be defined by genes related to cell size and structure, compared to other human neurons and mouse motoneurons? It is well established that human motoneurons are large, but to answer these relative size questions, we analyzed neuron soma size across all laminae in human and mouse lumbar spinal cord tissue. Given the obvious differences in overall body size and anatomy, we expected that most classes of human neurons would be larger than mouse neurons. Surprisingly, we found that, overall, human and mouse lumbar spinal neurons were approximately the same size, with a median Feret diameter (maximal caliper length) of 16.02 and 13.13 µm, respectively (human mean 20.3 ± 0.28 s.e.m; mouse mean 14.28 ± 0.12 s.e.m.) (Fig. 6C and Data File Table S2). Indeed, across most laminae of the spinal cord, human and mouse neurons displayed somewhat similar size distributions. By contrast, human lamina IX spinal neurons were approximately 2-fold larger than those in mouse and could be up to ∼120 µm across compared to ∼50 µm in mouse (Fig. 6E-F and Data File S5). These measurements are consistent with those previously reported for human and mouse spinal motoneuron soma (Ishihara et al., 2001; Kawamura and Dyck, 1977; McHanwell and Biscoe, 1981) and the same proportion that has been observed for human and mouse motoneuron axon caliber (M. D. Nguyen et al., 2000; Sobue et al., 1981a; 1981b). Assuming that human alpha motoneurons are within the higher end of this size distribution, then they are (1) much larger than other human spinal neurons, (2) increased in scale relative to mouse motoneurons, and (3) among the among the largest vertebrate neurons, including elephant motoneurons (∼85 µm) (Hardesty, 1902), human Betz corticospinal neurons (∼60-100 µm) (H. Braak and E. Braak, 1976), subsets of human dorsal root ganglion neurons (up to 100 µm) (Haberberger et al., 2019) and salmon Mauthner cells (∼87 µm) (Zottoli, 1978). This notable size of human motoneurons may explain the specialized gene expression signature that we observed in this subclass of neurons.

To assess specific ALS-related gene expression in tissue and to compare protein expression and *in situ* cell size, we next analyzed the protein expression of six ALS-related genes in post-mortem lumbar spinal cord from four donors, using immunofluorescence: NEFH, OPTN, PRPH, SOD1, STMN2, and TUBA4A. We found that neurons expressing NEFH, OPTN, PRPH, STMN2, and TUBA4A proteins were all enriched within the motoneuron region (lamina IX) of the lumbar spinal cord, with limited positive cells in other regions except for scattered, large cells in lamina III/IV of the dorsal horn which may be projection neurons and smaller neurons in medial lamina VII (Fig. 7A-D and Data File Table S5). SOD1 was present in lamina IX and throughout the spinal cord in a distinct peri-nuclear distribution, in contrast to the enriched RNA expression that we detected by single nucleus RNA sequencing. To ensure the accuracy of the SOD1 expression pattern, we validated the SOD1 antibody through targeted knockdown in human iPS neurons (Supplemental Fig. S18D). Overall, these data confirm the expression of ALS-related proteins in human spinal motoneurons in tissue (Pardo et al., 1995; Tsang et al., 2000).

We also studied the expression of these proteins in the mouse spinal cord, using lumbar tissue from aged animals (11 months old) to approximate the advanced age of the human subjects in this study. We found that Nefh, Optn, Prph, Stmn2, and Tuba4a displayed enrichment in lamina IX, while Sod1 was expressed ubiquitously, similar to what has been previously described for Sod1 in mice (Supplemental Fig. S18 A-C) (Pardo et al., 1995). Together with the comparative transcriptomic analysis above, this suggests that while human and mouse motoneurons are both enriched for expression of ALS-related genes, in human motoneurons the relative expression levels are higher and the enrichment of these genes as motoneuron-specific markers is greater.

Finally, we tested the relationship between expression of ALS-related genes and cell size within human spinal neurons in tissue. We measured the Feret distances of human neurons expressing each ALS-related protein in comparison with non-expressing neurons. We found that neurons that expressed NEFH, OPTN, PRPH, STMN2, and TUBA4A were generally larger than non-expressing neurons, both within the motoneuron region of lamina IX and in other lamina (Fig. 7D). Within lamina IX, this likely reflects enrichment within the larger alpha motoneurons (versus gamma) and in other laminae, this may reflect expression within spinocerebellar projection neurons that degenerate in ALS (Averback and Crocker, 1982; Williams et al., 1990) or other large cell classes. Importantly, we found that the very largest lamina IX neurons – that are known to be most susceptible to degeneration in ALS (Kawamura et al., 1981; McIlwain, 1991; Sobue et al., 1981b; 1981a) – were the most likely to express these markers. For lamina IX neurons with a Feret distance greater than 70 µm, on average 100% expressed NEFH, 81% expressed OPTN, 88% expressed PRPH, 60% expressed SOD1, 90% expressed STMN2, and 95% expressed TUBA4A (Data File Table S5).

These data further link motoneuron size and vulnerability to these cytoskeletal genes that have causative roles in motoneuron size and human disease.

## Discussion

The advent of single cell transcriptomic profiling approaches has transformed many aspects of biology and has the potential to pinpoint novel therapeutic targets amidst the complexity of human disease. In the context of the human nervous system in health and pathological conditions, multiple single nucleus RNA-sequencing studies of the cortex have been generated, but we still lack a comprehensive human spinal cord characterization that could provide crucial insights into chronic pain, spinal cord injury, and neurodegeneration. Here, we used human tissue samples prepared under careful neuroprotective conditions to collect high-quality tissue from organ donor subjects to create a cellular taxonomy of the adult human lumbar spinal cord including an atlas of glial, vascular, and neuronal cell types. We characterized the highly complex landscape of human neuronal signatures and contextualized these findings relative to established cell types in the mouse spinal cord. We revealed spatial location (along the dorsal-ventral axis) as the conserved, core organizing principle of mammalian spinal neurons. In addition, as a demonstration of the utility of this resource, we identified a signature of degeneration-associated genes expressed specifically in human motoneurons. This atlas and an accompanying web-based resource (https://vmenon.shinyapps.io/hsc_biorxiv/) serve as effective tools for understanding human spinal cord biology and enabling future discoveries.

This work builds on recent efforts toward understanding molecular and cellular heterogeneity in the human spinal cord, particularly during development. Rayon and colleagues (Rayon et al., 2021), focused on first trimester spinal cord derived from four human embryos, and identified diverse progenitor and neuronal populations, and performed a systematic comparison with the spinal cord cell types of the developing mouse spinal cord. Zhang and colleagues (Q. Zhang et al., 2021) profiled the early and mid-stages of fetal development with an important focus on glial development and cell-cell communication. For the adult human spinal cord, Zhang and colleagues performed single nucleus RNA-seq on the spinal cord from two donors and identified coarse glial and neuronal cell types (D. Zhang et al., 2021). However, they did not characterize human neurons to the same degree as this study, especially with respect to motoneurons, nor did they validate predicted gene expression patterns in tissue or provide a web resource for researchers to interact with the data.

Comparison of the human neuronal populations that we described here to their mouse counterparts can drive two major advances in our understanding of human spinal cord biology. First, the alignment of human cell types to mouse homologues for which the behavioral contributions and circuit information is available will allow us to extrapolate the function of human cell types. Together with overlaid human molecular data on disease markers or pharmacological responsiveness, these data will become a powerful perspective on pathophysiological mechanisms. Second, the discovery of conserved trends can identify core principles of spinal cord function. We previously found that dorsal and ventral neuronal populations displayed very different properties in their robustness, relatedness, and overall gene expression correlations. In addition, recent work that compared the influence of location, neurotransmitter status, and birthdate on perinatal spinal cell types also demonstrated the dominance of dorsal-ventral location in explaining spinal neuron variability. Together with these prior studies in mouse, our work here revealed that dorsal-ventral location is the shared, fundamental axis of spinal neuron transcriptional diversity.

Although we captured all major cell types and most known subclasses of cells in this catalog, we foresee further advances as additional data sets of this type are generated. In particular, the motoneuron population did not segregate into discrete subgroups based on molecular profile. This limitation may be technical, due to the overall signal-to-noise ratio of single-nucleus RNA-seq in key genes within prospective subgroups, or it may be that motoneuron substructure in adult human spinal cord is continuous at the transcriptomic level; studies of cortical and thalamic neurons have suggested the existence of such continuous transcriptomic variation(Bakken et al., 2021b; Tasic et al., 2018). As technological advances allow for higher-sensitivity transcriptomics on large numbers of cells, a clearer picture of the heterogeneity within motoneurons will likely become apparent. The current limitations did not affect our ability to identify robust signals distinguishing motoneurons from other classes of spinal neurons, especially when combining single-nucleus transcriptomics with spatial approaches. Rather, this resource allowed us to observe key aspects of the human motoneuron expression profile that support a model of specific molecular repertoires for motoneuron cell structure that also confer selective vulnerability to degeneration(Castellanos-Montiel et al., 2020; Clark et al., 2016; Hardy and Rogaeva, 2014).

An intriguing finding from our analysis, made possible by our extensive profiling of motoneurons, is the enrichment of cytoskeletal gene expression in these cells. All cells require a functional cytoskeleton, raising the question of why spinal motoneurons in particular are so crucially dependent on the proper expression and function of cytoskeletal-related genes. Interestingly, the neurofilament genes that were enriched in human spinal motoneurons compared with other neuronal populations – NEFL, NEFM, NEFH, and PRPH – are precisely those structural components that drive increased axon caliber and cell size (Friede and Samorajski, 1970; Hoffman et al., 1984; Lee and Cleveland, 1996; M. D. Nguyen et al., 2000). Over-expression of mouse NEFL, human NEFM, human NEFH, or mouse PRPH in transgenic mice can each cause enlargement and swellings of motoneuron somas and subsequent axon degeneration (Beaulieu et al., 1999; Côté et al., 1993; Gama Sosa et al., 2003; Marszalek et al., 1996; Xu et al., 1993b; 1993a), linking human motoneuron gene expression and cellular phenotype. Relatedly, these neurofilament genes are found in other large neurons in the brain and peripheral nervous system, suggesting that they may be part of a common signature that permits increased cell size (Bakken et al., 2021a; Limone et al., 2021; M. Q. Nguyen et al., 2021; Tsang et al., 2000; Zeisel et al., 2018). Large soma size and axon caliber may be required to sustain extensive dendritic trees and axons up to a meter long, to support cell energetics, or for firing rate and conduction parameters (Manuel et al., 2019; Perge et al., 2012; Schoenen, 1982). These large cells then rely critically on this protein network and are selectively vulnerable to its abnormal function. Human motoneurons were also distinguished by expression of the microtubule stability factors TUBA4A and STMN2 (Clark et al., 2016; Klim et al., 2019), potentially highlighting a requirement for structural support in these peripherally projecting cells subject to axonal wear and tear during body movement.

Finally, it is critical to consider the spinal cord as a *community* of cell types that function together in normal health and disease. While we highlight the molecular signature of motoneurons, the single nucleus RNA-sequencing data set that we present provides the first comprehensive resource of all cell types in the adult human spinal cord. We anticipate that this work will have broad implications for understanding spinal cord biology, allowing researchers to parse how ubiquitous genetic alterations interact with diverse cell-type specific molecular profiles in disease and how particular populations may respond to target molecular interventions and pharmacology in chronic pain. We hope that our work will serve as a broad resource and foundation for studying the wide range of cell types involved in sensory and motor function in the human spinal cord.

## STAR★Methods

### Key resources table

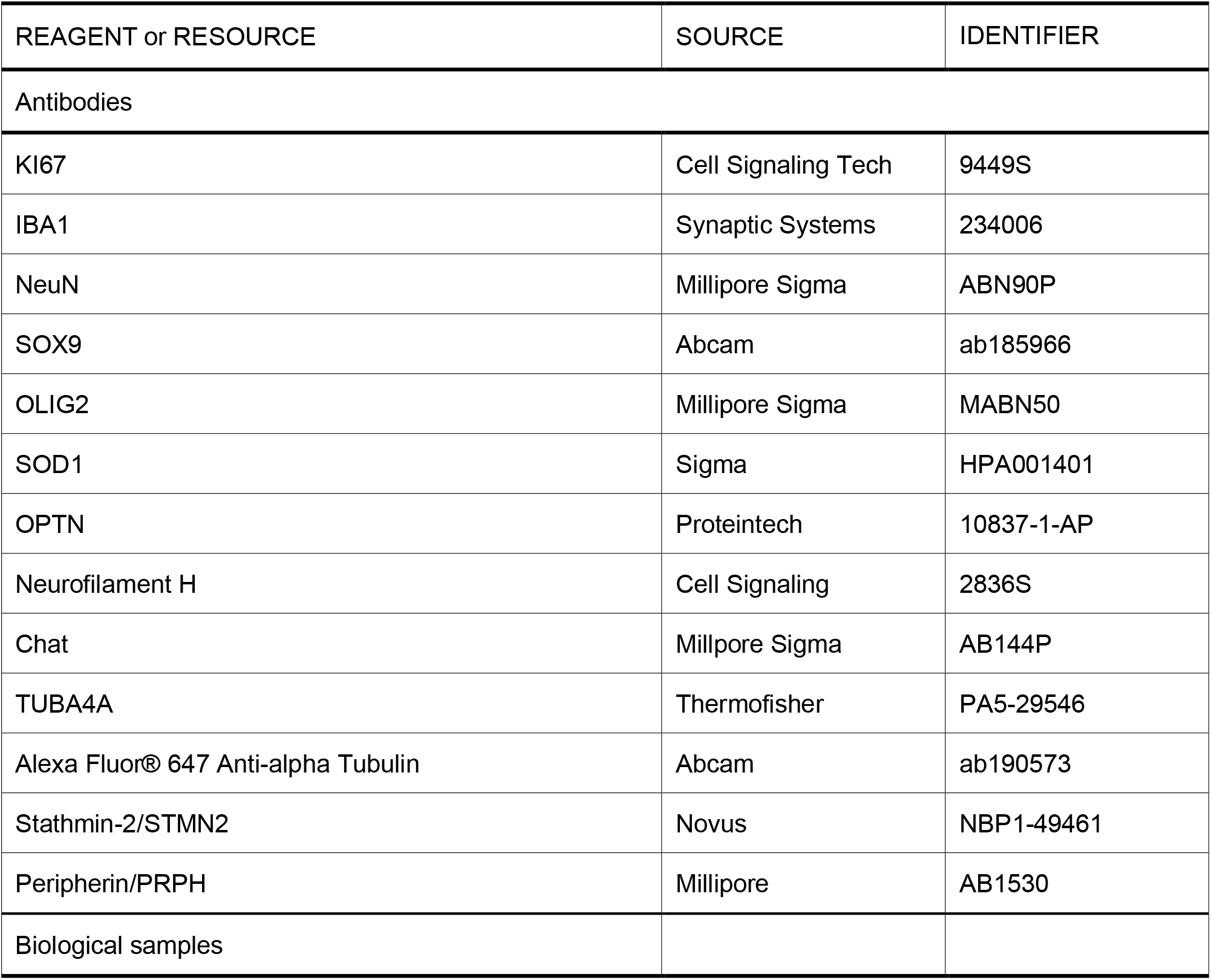
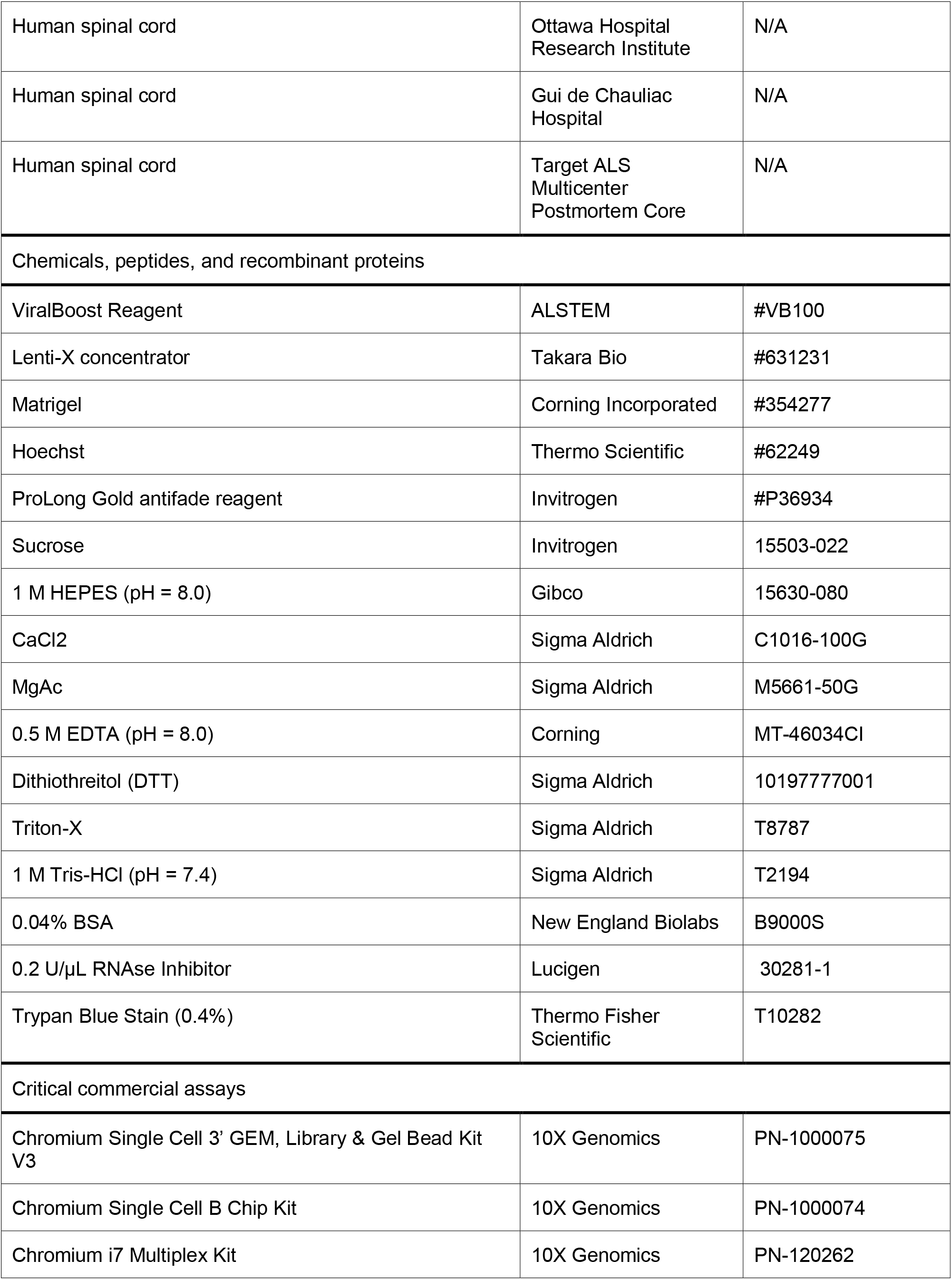
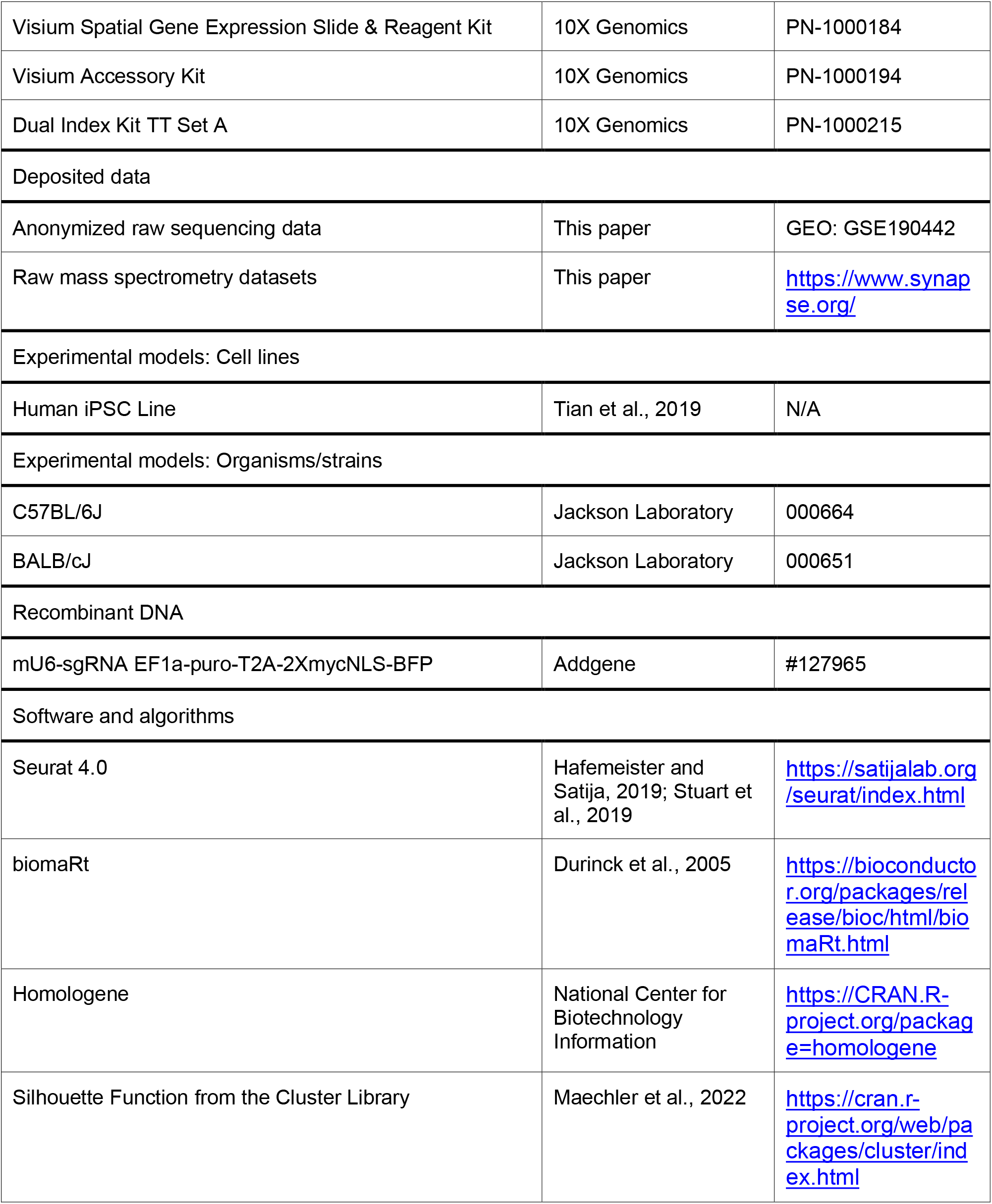
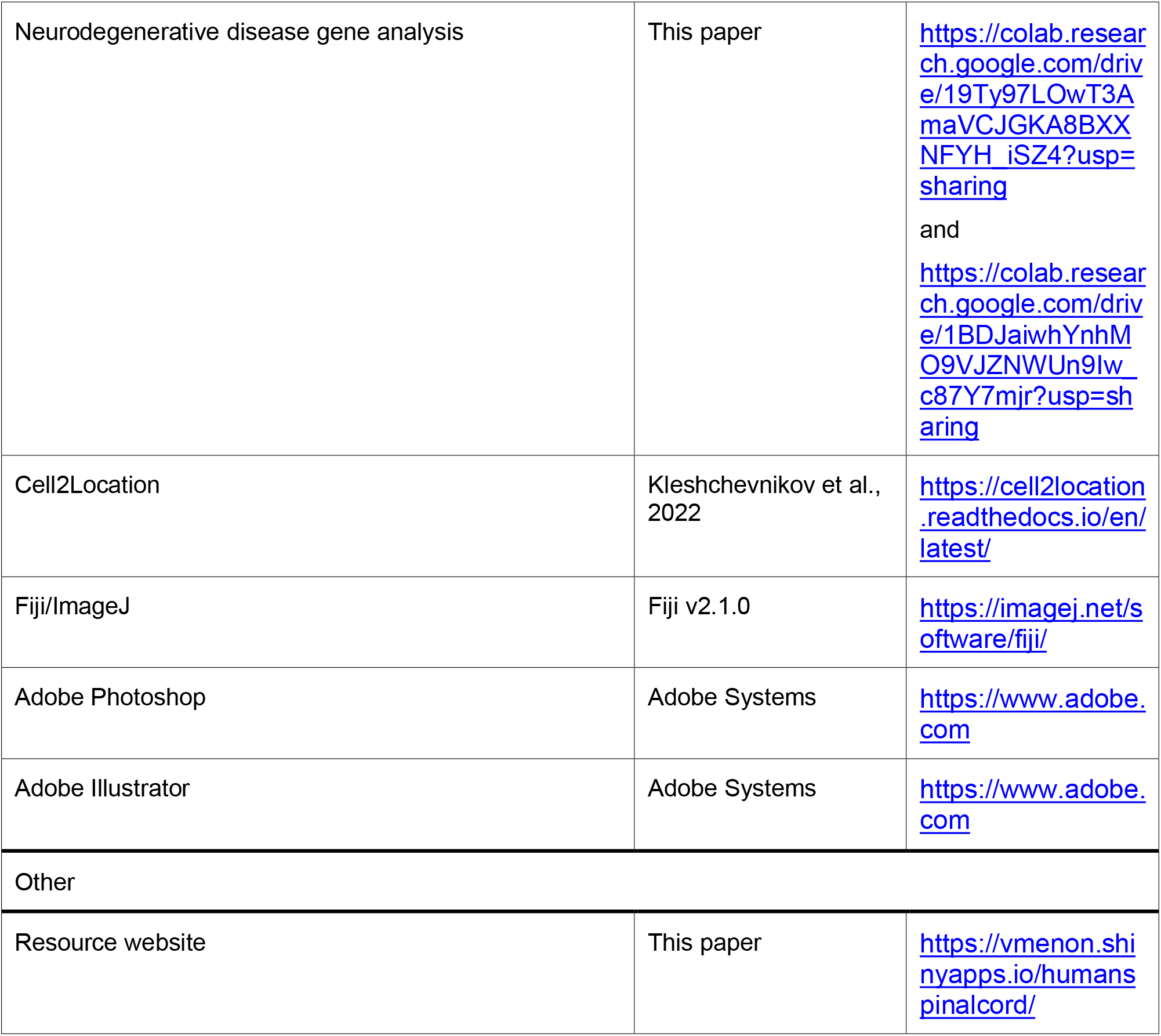

#### Human spinal cord acquisition and preparation

Spinal cords for single nucleus RNA sequencing were obtained from neurologic determination of death organ-donor patients (∼50-80 years old, 4 men, 3 women) under the approval of the French institution for organ transplantation (Agence de la Biomédecine) or the Ottawa Health Science Network Research Ethics Board, following the template provided by the University of Ottawa and the Tri-Council Policy Statement Guidelines. Both approvals imply consent for using anonymized donor genetic information. Human lumbar spinal cords were retrieved under chilled body and neuroprotective conditions as described previously (Bauchet et al., 2022; Dedek et al., 2019; Galuta et al., 2020). The extraction procedure took 20-40 minutes and was done within three hours of cessation of circulation by aortic cross-clamp. Lumbar spinal cord tissue was flash frozen on liquid nitrogen in the operating room and stored at −80°C until nuclei isolation.

For immunohistochemistry experiments, lumbar spinal cord tissue was isolated from organ-donor patients (∼55-65 years old, 3 men, 1 woman). The tissue was immediately fixed in 4% paraformaldehyde for 24-48 hours, then washed in PBS, and placed in 30% sucrose for 2-4 days at 4°C before being embedded in OCT medium for sectioning.

For Visium spatial transcriptomics, postmortem lumbar spinal cord from a non-neurological control subject (∼75 years old, male) was acquired from the Target ALS Multicenter Postmortem Core as part of the New York Genome Center (NYGC) Amyotrophic Lateral Sclerosis (ALS) Consortium. Informed consent is acquired by each Target ALS member site through its own institutional review board (IRB) protocol and samples are transferred to the NYGC in accordance with all applicable foreign, domestic, federal, state, and local laws and regulations for processing, sequencing and analysis. The Biomedical Research Alliance of New York (BRANY) IRB serves as the central ethics oversight body for the NYGC ALS Consortium. Ethical approval for this study was given by the BRANY IRB.

#### Mouse work and spinal cord acquisition

All procedures and experiments were approved by the Animal Care and Use Committee of NINDS (protocol #1384). Adult mice were of 50:50 mixed background from strains C57BL/6J and BALB/CJ, housed in standard conditions. For basic anatomical experiments, two male and two female mice of approximately 24 weeks old were used. For ALS marker gene expression studies, two male and one female mice of approximately 11 months old were used. To obtain spinal cord tissue, anesthetized mice were transcardially perfused with PBS followed by cold 4% paraformaldehyde (PFA). The spinal cords were harvested and post-fixed in cold 4% PFA overnight at 4°C, cryoprotected by immersion in 30% sucrose overnight at 4°C and embedded in OCT medium for sectioning.

#### Nuclei isolation

Nuclei were isolated from fresh frozen human spinal cords using a triton-based protocol (Matson et al., 2018). Briefly, after removing the dura, half a segment of spinal cord was placed in a Dounce homogenizer (Kontes Dounce Tissue Grinder) containing 500 μL of lysis buffer (0.32 M sucrose, 10 mM HEPES [pH 8.0], 5 mM CaCl2, 3 mM 586 MgAc, 0.1 mM ETDA, 1 mM DTT, 0.1% Triton X-100). After douncing with 5 strokes of pestle A and 5-10 strokes of pestle B, the lysate was diluted in 3 mL of sucrose buffer (0.32 M sucrose, 10 mM 588 HEPES [pH 8.0], 5 mM CaCl2, 3 mM MgAc, 0.1 mM ETDA, 1 mM DTT) and passed over a 70 μm strainer. The filtered lysate was centrifuged at 3,200 x g for 5 min at 4°C. After centrifugation, the pellet was resuspended in 3 mL of sucrose buffer and centrifuged again at 3,200 x g for 5 min at 4°C. After centrifugation, the pellet was resuspended in 3 mL sucrose buffer and incubated for 2 min on ice. The sample was transferred to an Oak Ridge tube and homogenized for 1 min using an Ultra-Turrax Homogenizer (IKA). Then, 12.5 mL of density sucrose buffer (1 M sucrose, 10 mM HEPES [pH 8.0], 3 mM MgAc, 1 mM DTT) was layered below the sample. The tube was centrifuged at 3,200 x g for 20 min and the supernatant immediately poured off. The nuclei on the side of the tube were resuspended with 100 µL of PBS with 0.04% BSA and 0.2 U/µL RNase inhibitor. Nuclei were inspected for visual appearance and quantified with a hemocytometer before proceeding with nuclei capture and sequencing.

#### Single nucleus RNA sequencing

Single nucleus RNA sequencing was carried out using Single-cell gene expression 3’ v3 kit on the Chromium platform (10X Genomics) according to manufacturer’s instructions with one modification. Following reverse-transcription, an additional PCR cycle was added to the number of cycles for cDNA amplification to compensate for decreased cDNA abundance in nuclei compared to cells. Libraries were sequenced to a minimum depth of 20,000 reads per nucleus using an Illumina HiSeq 3000 (PE 26 – 8 – 98 bp). Raw sequencing reads were demultiplexed, aligned, and a count matrix was generated using CellRanger. For alignment, introns and exons were included in the reference genome (GRCh38).

#### Quality check analysis

All the 10x runs for each human sample were initially filtered with an nUMI cutoff of >1000 and then nuclei with less than 5% mitochondrial gene contamination were retained. Next, the mitochondrial genes were also removed from the matrices.

#### Top level UMAP and clustering

The 8 human datasets were integrated using SCTransform normalization followed by CCA based integration from Seurat 4.0 (Hafemeister and Satija, 2019; Stuart et al., 2019).

The integrated sets were then jointly analyzed to identify optimal Principal Component values based on ElbowPlot and PCheatmaps. PC value of 30 was used for clustering and UMAP. The clusters, obtained using a value of 0.6 for Seurat’s resolution parameter, were then manually annotated based on the expression of marker genes for various cell types, namely neurons, astrocytes, microglia, oligodendrocytes, OPCs, endothelial cells, pericytes, meningeal cells, Schwann cells, and lymphocytes.

#### Sub clustering of major cell types

Identification of subclusters within cell types was performed separately for three major cell types (neurons, microglia, astrocytes), with the rest being subclustered as groups (Group 1-oligodendrocytes, OPCs, and Schwann cells; Group 2-endothelial cells, pericytes, meningeal cells, and lymphocytes). For each cell type/group, the subclustering was done in multiple rounds until no putative transcriptomic doublets or contamination of other cell types was observed (described below).

For subclustering of major cell types, the raw counts were extracted from 8 datasets, for each cell type, and then re-normalized (using log normalization) and scaled in order to prepare for integration. The integration of 8 datasets belonging to a particular cell type was performed based on CCA-integration workflow from the Seurat 4.0 package. Optimal PC values were selected based on ElbowPlot and PCheatmaps for each cell type in order to be used for further sub clustering and preparation of UMAPs. Multiple resolutions were interrogated, depending on cell type, ranging from values of 0.08 to 3.

During each round, putative transcriptomic doublet clusters and contamination of other cell types was removed (based on expression of multiple major class genes) and the above steps were performed again. Doublets were identified by clusters that expressed markers for more than one cell type. All clusters were checked for doublets by their markers using wilcox and auroc tests, as well as visually using the FeatureScatter option in Seurat.

#### Subclustering of neurons

Neurons were clustered in 2 stages, first dividing the neurons into motoneurons, ventral neurons and dorsal neurons, followed by a second round of further subclustering within motoneurons, ventral neurons and dorsal neurons. As described in the main text, dorsal and ventral neuronal groups were identified using marker genes from previous studies on mouse neurons.

During the first stage, the log normalization of raw counts and scaling (including regressing out the number of transcripts and mitochondrial percentage) of each dataset was done followed by integration based on the same steps described above for the glia and vascular cells. During the second stage, raw counts were again extracted from each group (motoneurons, ventral and dorsal) and normalized using SCTransform (to avoid dataset size related limitations) and followed by the standard integration workflow in the Seurat 4.0 package. In order to obtain a refined set of neuronal subpopulations, all the subclusters were interrogated for ‘low quality’ (based on gene detection), and doublets and other contamination and were subsequently removed from the analysis. All the refined clusters were then re-integrated to prepare a combined neuronal UMAP and mapped with refined subcluster annotations.

During each subclustering, cluster-specific genes were identified based on Wilcox Rank-Sum test and ROC analysis within the FindMarkers function from Seurat 4.0. Based on these genes, the distinct subpopulations based on expression of candidate markers were manually annotated.

#### Cluster robustness assessment and silhouette scores

We used two approaches to assess cluster robustness: a post-hoc machine learning-based classification approach, and a silhouette score approach.

For the post-hoc machine learning approach, we built a random forest classifier for every pair of neuronal clusters, trained on 80% of the nuclei. This classifier was then used to assign cluster membership for the remaining 20% of the cells, and the entire process repeated such that each cell in every pairwise cluster comparison was classified 100 times. A cell that was classified into its original cluster <90 times was deemed “misclassified”. For every pair of clusters, we then calculated the mean percentage of cells that were misclassified among the two clusters to generate pairwise cluster robustness scores. For visualization as a constellation diagram, we only connected cluster pairs with minimum misclassification percentage >3%, representing their connections with the mean misclassification percentage.

For silhouette score evaluation, we used the ‘silhouette’ function from the ‘cluster’ library in R (https://cran.r-project.org/web/packages/cluster/index.html), where the Euclidian distance matrix based on the first 25 PCs was used as input, together with the neuronal cell type annotations.

#### Tissue processing, Visium data generation, and Visium data preprocessing

Frozen postmortem lumbar spinal cord from a non-neurological control subject was embedded in Tissue Plus OCT Compound (Fisher Healthcare, catalog no. 4585) and cryosectioned at −16°C. Sections of 10 µm thickness were collected onto prechilled Visium Spatial Gene Expression Slides (10x Genomics, catalog no. 1000185) by warming the back of the slide to adhere the tissue. Visium spatially resolved gene expression data was generated according to the Visium Spatial Gene Expression User Guide (10x Genomics, CG000239 Rev F). Briefly, tissue sections were fixed in chilled methanol and stained using hematoxylin and eosin. Brightfield histological images were acquired using an EC Plan-Neofluar 10x/0.3 M27 objective on a Zeiss Axio Observer Z1 fitted with a Zeiss Axiocam 506 mono (Carl Zeiss Microscopy, Germany). Raw CZI images were stitched using Zen 2012 (blue edition) (Carl Zeiss Microscopy, Germany) and exported as JPEGs. Tissue sections were permeabilized for 12 minutes which was selected as the optimal time based on tissue permeabilization time course experiments conducted using the Visium tissue optimization protocol. cDNA libraries were prepared and quantified according to the Visium Spatial Gene Expression User Guide (10x Genomics, CG000239 Rev F) and pooled at a concentration of 10 nM for sequencing.

Pooled spatial gene expression libraries were loaded at a concentration of 0.9 nM and sequenced on a NovaSeq 6000 System using a NovaSeq S4 Regent Kit v1.5 (200 cycles, Illumina, catalog no. 20027466) using the following recipe: read 1: 100 reads, i7 index read: 10 cycles, i5 index read: 10 cycles, read 2: 100 cycles. The average sequencing depth for each sample was approximately 200-280 x 106 reads.

Raw FASTQ files and histological images were processed using Space Ranger v.1.3.0, which uses a modified STAR v2.7.2a for genome alignment and performs barcode/UMI counting to generate feature-spot matrices. Reads were aligned to a GRCh38 reference genome filtered to exclude lncRNAs, pseudogenes and mitochondrially encoded genes.

#### Cross species analysis between human spinal cord vs mouse meta-analysis datasets

Cross species comparison between human and mouse meta-analysis (Russ et al., 2021) spinal cord datasets were performed at two levels: 1. “Top level” which includes all major cell types and 2. Neurons only.

In both cases, the orthologous genes within mouse data matrix were converted to human homologs using biomaRt package (Durinck et al., 2005) from Bioconductor and in-house scripts. The raw counts from both human and mouse datasets were then split by different samples and then re-normalized, scaled and integrated. For the “top-level” analysis, SCTransform based integration was performed whereas for neurons only, log normalization-based integration was performed. Subsequently, UMAPs and correlation matrices were generated for further cross-species comparison of various cell types at top level and neuronal sub-clusters.

#### Cross-correlation of human and mouse cluster expression

Cross-species cluster correlation measures were calculated from PCs in the integrated space (using 20 PCs for the top level comparison of major cell classes, and Pearson’s correlation of the top 2,000 highly variable genes. Aggregate correlation values for each pair of clusters (one mouse, one human) were calculated as the mean correlation value across all human-mouse nuclei pairs from the respective clusters.

Quotient graphs using qgraph in R were used to show the correlations greater than 0.8 based on the top 2,000 highly variable genes between human and mouse spinal cord neurons (graph “cor”, layout “spring”).

#### GO analysis of human motoneuron marker genes

The top markers (based on smallest adjusted p-value) of human spinal motoneurons were determined based on the Wilcox Rank Sum test and were analyzed using DAVID 6.8 GO enrichment analysis (https://david.ncifcrf.gov/summary.jsp). The general categories of GOTERM_BP_DIRECT, GOTERM_CC_DIRECT, and GOTERM_MF_DIRECT were analyzed and functional annotation clustering was performed using default parameters including medium classification stringency.

#### Focused comparison of mouse and human motor neurons

Human motor neurons were compared to mouse lumbar skeletal motor neurons from a recent study (Alkaslasi et al., 2021). Mouse MN genes were converted to human homologs using Homologene (https://CRAN.R-project.org/package=homologene). Only genes with human homologs present in both datasets were included in the analysis (13,574). Raw counts were extracted from each original dataset, normalized using SCTransform, and integrated based on integration anchors. Clustering was performed as described above (resolution = 0.4), and differentially expressed genes were identified based on Wilcox Rank Sum test and ROC analysis within FindMarkers function from Seurat 4.0.

#### Analysis of evolutionarily convergence/divergence scores

All available data on gene expression-based human:mouse divergence scores was downloaded from Pembroke et al (Pembroke et al., 2021). Genes of interest were then extracted, yielding scores for three genes (SOD1, TUBA4A, OPTN) that overlapped with this data. We compared the mean and standard deviation of these three genes to the same metrics for the remainder of the assayed genes from the Pembroke report (N=1426 other genes) using a standard two-sided t-test.

#### Neurodegenerative disease gene analysis

Post-QC scRNAseq count data was extracted for seven major cell classes of interest. For each gene per cell class, mean expression was calculated across all assayed cells of that class. These means were then transformed to a Z scale to facilitate comparisons across multiple cell types. The Z scaling was carried out using the mean and standard deviations as the scaling functions as this is the common convention for this conversion. Additionally, genes that did not have any count based data available for that cell class were set to zero at Z scaling. From this large dataframe of normalized counts per cell type, candidate genes for HSP, PD and ALS were extracted from Genomics England Expert Panel App genes audited at the “green” level of confidence [https://panelapp.genomicsengland.co.uk/]. AD genes were annotated by an expert panel and extracted (Ramos et al., 2021) and the ALS list was also supplemented with genes from the literature, as described in the main text. These extracted genes were then clustermapped using the python package seaborn with Z scores greater than 7 truncated to a value of 7 for display purposes.

#### SOD1 antibody validation in human iPS neurons with targeted knockdown

Previously published human inducible pluripotent stem cells (hiPSCs) were used to knock down SOD1 (Tian et al., 2019). A SOD1 or non-targeting control sgRNA was cloned into a mU6-sgRNA EF1a-puro-T2A-2XmycNLS-BFP vector (gift from Martin Kampmann’s lab; Addgene #127965). sgRNA sequences are as follows: SOD1: GAGGCACCACGACAGACCCG, non-targeting sgRNA: GAATATGTGCGTGCATGAAG. Lentivirus was produced via transduction of Lenti-X HEK 293T cells using Lipofectamine 3000 in DMEM high glucose GlutaMAX Supplement media containing 10% FBS. 24 hours post-transfection, media was replaced, including ViralBoost Reagent (ALSTEM, #VB100). 96 hours post-transfection, media was collected and concentrated 1:10 in 1xPBS using Lenti-X concentrator (Takara Bio, #631231), aliquoted, and stored at –80°C. 100 ml of these aliquots was used to transduce 100,000 hiPSCs to generate SOD1 KD and control lines. The cells were split and replated on Matrigel (Corning Incorporated #354277) coated coverslips with the viral concentrate in E8+Y-27632 ROCK Inhibitor and allowed to incubate for 24 hours at 37°C, 5% CO_2_. The media was replaced with E8 and the cells were allowed to grow for another 24 hours before fixation with 4% PFA in PBS for 10 mins at room temperature. Cells were washed with PBS 3 times and permeabilized in block (PBS + 3% donkey serum + 0.1% tritonX) for 30 mins at room temperature. Primary antibody targeting SOD1 (Sigma, HPA001401-100UL) was diluted at 1:500 in block and cells were incubated in primary overnight at 4°C on a rocker. The next day, cells were washed three times with PBST and incubated in block with secondary antibody (Jackson ImmunoResearch # 711-625-152) and Hoechst (Thermo Scientific #62249) for 1 hour at room temperature. Following 3 washes with PBST, the coverslips were mounted using ProLong Gold antifade reagent (Invitrogen #P36934). After curing, the coverslips were imaged using Nikon spinning disk confocal using laser wavelengths of 405 nm, 488 nm, 561 nm, and 640 nm at 100ms exposure and 75%, 25%, 25% and 100% power respectively. Images were edited using ImageJ.

#### Immunohistochemistry antibodies

KI67 (Cell Signaling Tech, 9449S), IBA1 (Synaptic Systems, 234006), NeuN (Millipore Sigma, ABN90), SOX9 (Abcam, ab185966), OLIG2 (Millipore Sigma, MABN50), SOD1 (Sigma, HPA001401-100UL), OPTN (Proteintech, 10837-1-AP), Neurofilament H (Cell Signaling, 2836S), Chat (Millpore Sigma, AB144P), TUBA4A (Thermofisher, PA5-29546), Alexa Fluor® 647 Anti-alpha Tubulin (Abcam, ab190573), Stathmin-2/STMN2 (Novus, NBP1-49461), and Peripherin/PRPH (Millipore, AB1530).

#### Immunohistochemistry

Immunohistochemistry for human and mouse spinal cords were performed as previously described (Sathyamurthy et al., 2018) with modifications for human spinal cords. Briefly, mouse spinal cords were cut at 50 µm and blocking buffer (1% IgG-free BSA, 10% normal donkey serum, 0.1% Triton-X 100 in PBS) for one hour, prior to incubation in blocking buffer and primary antibody for 48 hours at 4°C. Primary antibody was washed off three times in PBS before a 2-hour incubation in secondary antibody at room temperature. Secondary antibody was washed off three times in PBS before adding a coverslip.

Human spinal cords were cut at 14 µm, washed twice in TBS and placed in 0.05% sodium azide-TBS at 4°C for 3 days under a LED light to quench autofluorescence. Human spinal cords were placed in blocking buffer (1% IgG-free BSA, 10% normal donkey serum, in TBS) for one hour prior to incubation in blocking buffer and primary antibody for 48 hours at 4°C. Primary antibody was washed off three times in TBS with 0.025% triton before a 2-hour incubation in secondary antibody at room temperature. Secondary antibody was washed off three times in TBS with 0.025% triton before adding a coverslip.

#### Imaging

Images of immunohistochemistry samples were imaged using a Zeiss 800 LSM confocal microscope.

#### Image analysis and quantification

The images were overlaid in Adobe Photoshop where borders between the gray and white matter and the lamina within the gray matter were drawn. These images were then exported to ImageJ for analysis. The cells were measured manually by outlining each cell using the selection tool and adding them to groups within the ROIManager in ImageJ based on lamina. Feret diameter measurements of all the ROIs for each section were saved in a spreadsheet. The white and gray matter of each subject were outlined in ImageJ and their areas were exported to a spreadsheet.

To identify colocalization of markers with NeuN, each neuron was first outlined with the selection tool in ImageJ and saved into different groups based on whether the cell was in lamina IX or not. Then, each cell that had co-occurrence of the markers were placed into separate groups (double positive in lamina IX and double positive outside lamina IX). Feret diameter measurements were then saved to a spreadsheet and the number of cells in each group were counted in Python.

#### Statistical testing

Two-way ANOVA (repeated-measures) was used for assessing grouped data, such as the correlation and silhouette scores between human and mouse dorsal vs ventral neurons. Two-tailed t tests (unpaired) were used all for differences in silhouette scores and correlation between clusters as well as expression of protein and soma size, as indicated in figure legends. Bonferroni-adjust Wilcox test p-values and Bhattacharyya Coefficients (BC) were used for comparison of human vs mouse cell diameter. Differences among groups were considered significant if p < 0.05. P values are denoted by asterisks: ∗p < 0.05; ∗∗p < 0.01; ∗∗∗p < 0.001; ****p < 0.0001; n.s – not significant. Data are represented as mean ± s.e.m. unless otherwise indicated. Statistical analyses were performed using GraphPad prism software and R.

#### Data and code availability

Anonymized raw sequencing data and counts tables are deposited in the Gene Expression Omnibus (GEO) with accession number GSE190442 and with associated metadata in Data File Table S7. The raw mass spectrometry datasets are deposited with Synapse.org. In addition, visualization of expression data at the cluster and donor level are available through a searchable web resource at https://vmenon.shinyapps.io/hsc_biorxiv/.

## Supplemental Materials

Figures. S1 to S21

Data Files Tables S1 to S8

## Acknowledgments

We gratefully acknowledge the gift of human tissue from the 11 donors included in this work and their families, whose contribution has been critical for this work. We thank Dr. Yue Andy Qi for assistance with protein validation. This work was supported by NINDS Intramural funds through 1ZIA NS003153 (KJEM, LL, IH, AJL), 1ZIANS003155 (SH and MEW) NS116350 (JP, KK, HP); by NICHD Intramural funds through 1ZIAHD008966 (MRA, CELP); by the Intramural Research Program of the NIA project ZO1 AG000535 (MAN); by R01 AG06683 and U54 AG076040 from NIA and the NIH Common Fund (AY, VM); by the Canadian Institutes of Health Research, the University of Ottawa Department of Surgery, the Ontario Neurotrauma Foundation, and the Praxis Spinal Cord Institute (AG, SA, JP, MMA, FAQ, SMA, APW, ECT, MEH); and by ANR-15-CE16-012, FRC-EET-2019 grants, CNRS/INSERM/Montpellier Hospital research supports (PFM, EB, LB) and a AL210154 grant (JP, KK, HP).

## Author contributions

Conceptualization: V.M., A.J.L.

Methodology and Investigation: all authors.

Formal Analysis: A.Y., K.J.E.M., I.H., M.R.A., M.A.N., V.M., A.J.L.

Visualization: A.Y., K.J.E.M., I.H., M.R.A., V.M., A.J.L.

Writing: K.J.E.M., M.E.W., C.E.L.P., V.M., A.J.L.

Supervision: M.E.W., M.E.H., P.F.M., E.B., L.B., E.C.T., H.P., C.E.L.P., V.M., A.J.L.

Funding acquisition: M.A.N., M.E.W., M.E.H., P.F.M., E.B., L.B., E.C.T., H.P., C.E.L.P., V.M., A.J.L.

## Supplemental Materials

**Supplemental Fig. S1.**
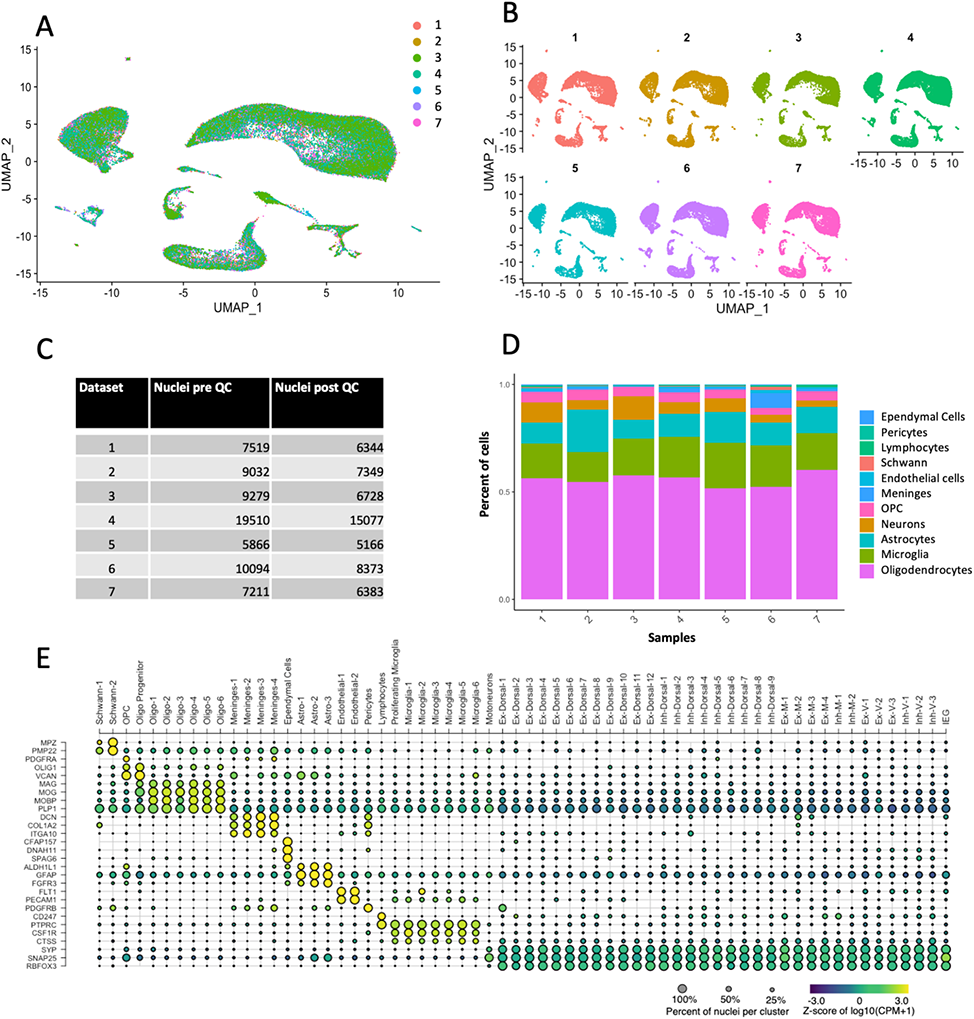
Integration at the top level and identification of major cell types. **A,** UMAP representation of the 55,420 nuclei after integration of the 7 human datasets. **B,** UMAP from panel A split by datasets to depict the overlap between datasets. **C,** Number of nuclei before and after quality check analysis (includes removal of doublets, low quality and other cell-type based contaminations). The number of nuclei in panel A are equal to total post QC. **D,** Bar plot showing the proportion of nuclei assigned as a particular cell type per dataset. **E,** Dot plot showing the expression of 28 canonical marker genes in all the major cell types and their subclusters (also depicted as UMAP representation in figure 1B in main manuscript. Microglia-1-5, Meninges-1-4, Endothelial-1-2 corresponds to Micro-1-5, Men-1-4, Endo1-2, respectively; in Fig 1B). Ex- Excitatory, Inh- Inhibitory, M- Mid, V- Ventral.

**Supplemental Fig. S2.**
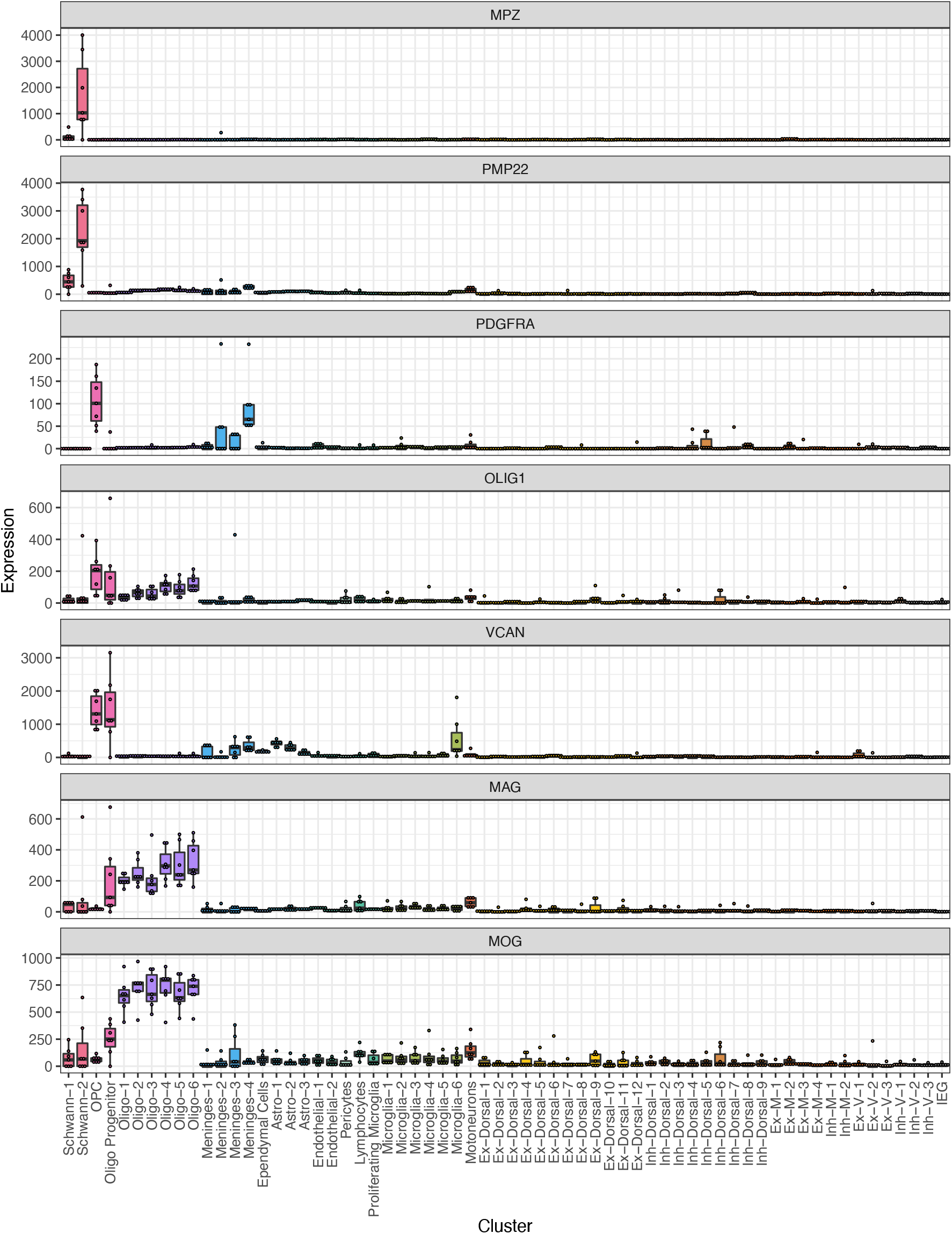
Expression of marker genes in all human spinal cord cell. Box plot representation of per-cluster and per-sample expression (Counts per Million) of top-level marker genes in all cell types. Ex- Excitatory, Inh- Inhibitory, MV-Mixed ventral.

**Supplemental Fig. S3.**
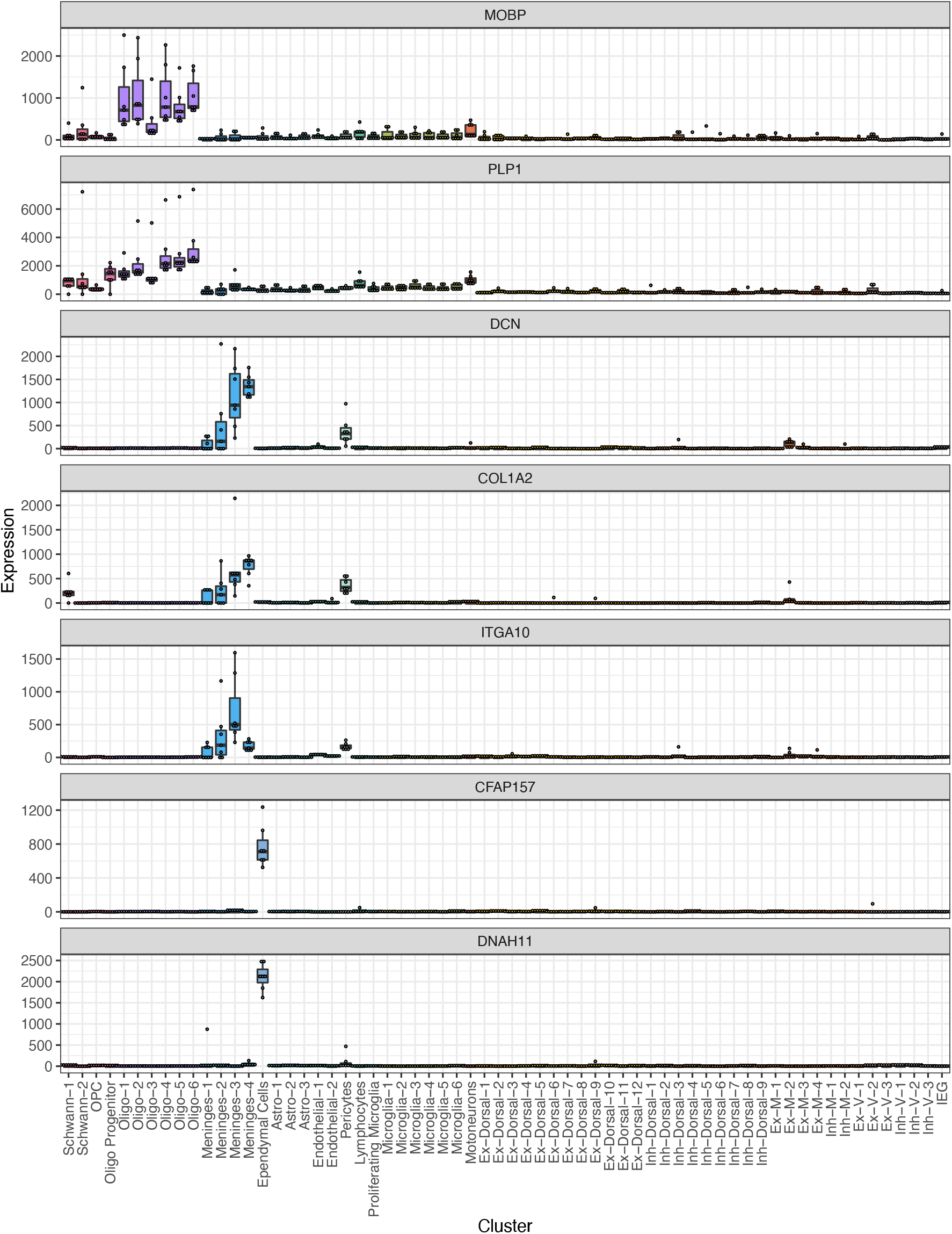
Expression of marker genes in all human spinal cord cell. Box plot representation of per-cluster and per-sample expression (Counts per Million) of top-level marker genes in all cell types.

**Supplemental Fig. S4.**
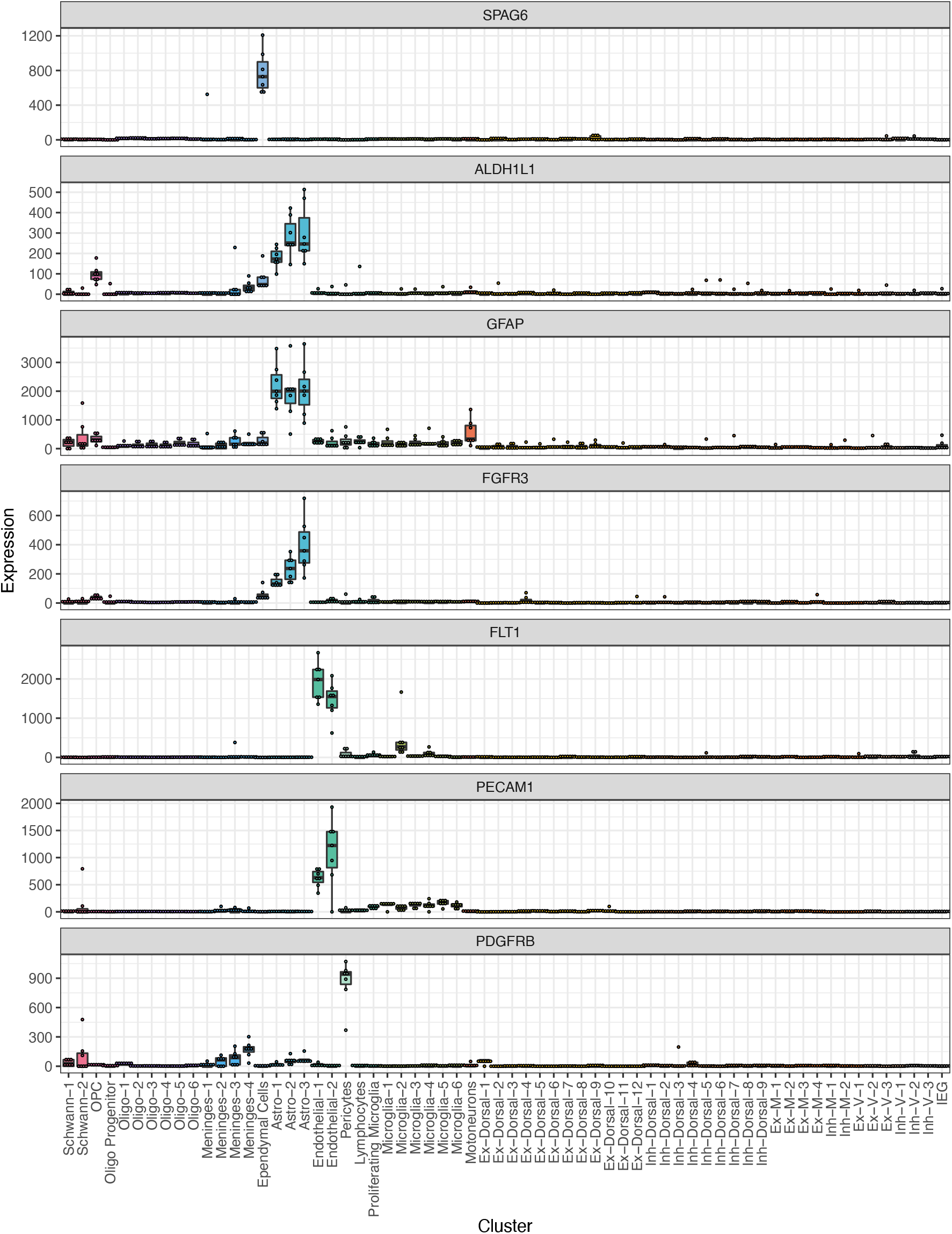
Expression of marker genes in all human spinal cord cell. Box plot representation of per-cluster and per-sample expression (Counts per Million) of top-level marker genes in all cell types.

**Supplemental Fig. S5.**
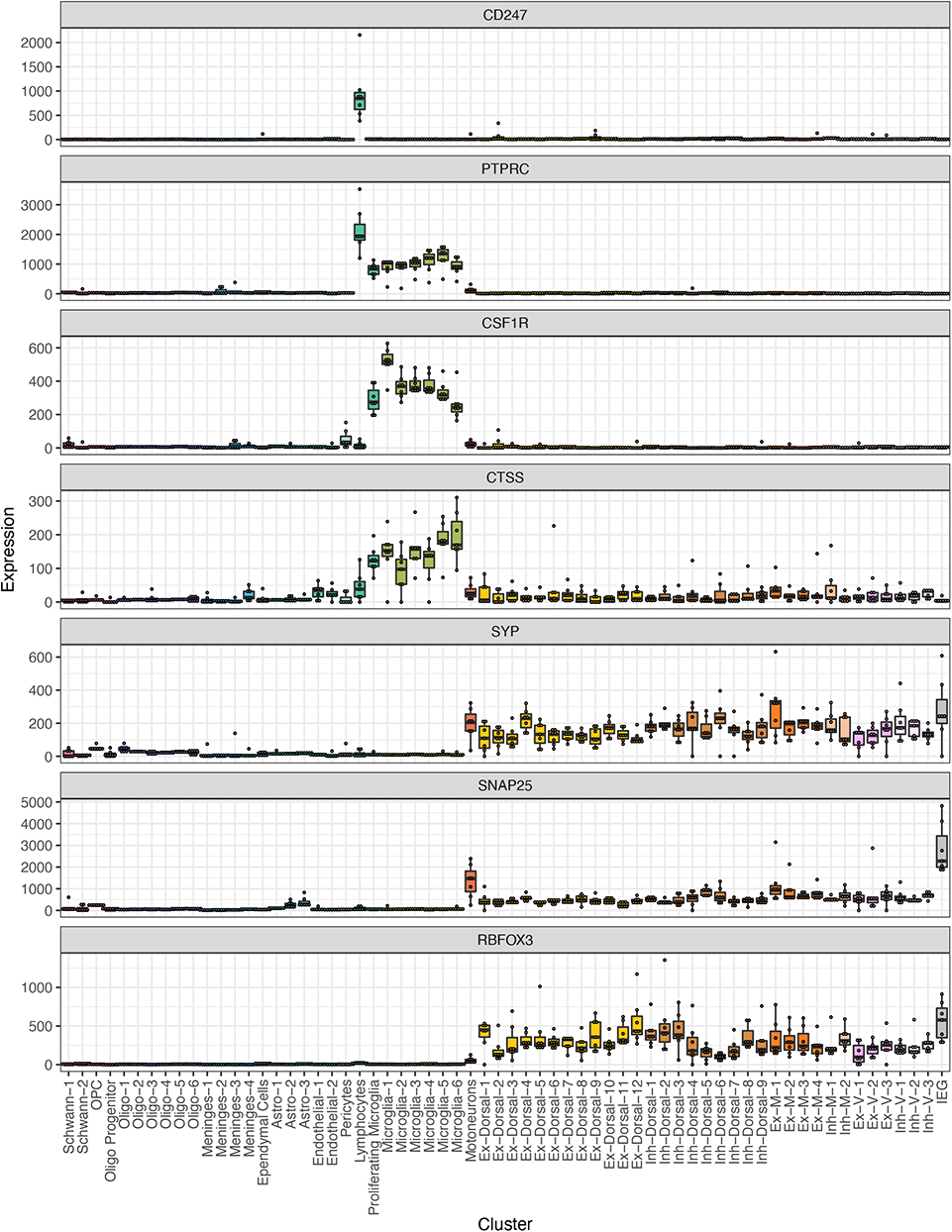
Expression of marker genes in all human spinal cord cell. Box plot representation of per-cluster and per-sample expression (Counts per Million) of top-level marker genes in all cell types.

**Supplemental Fig. S6.**
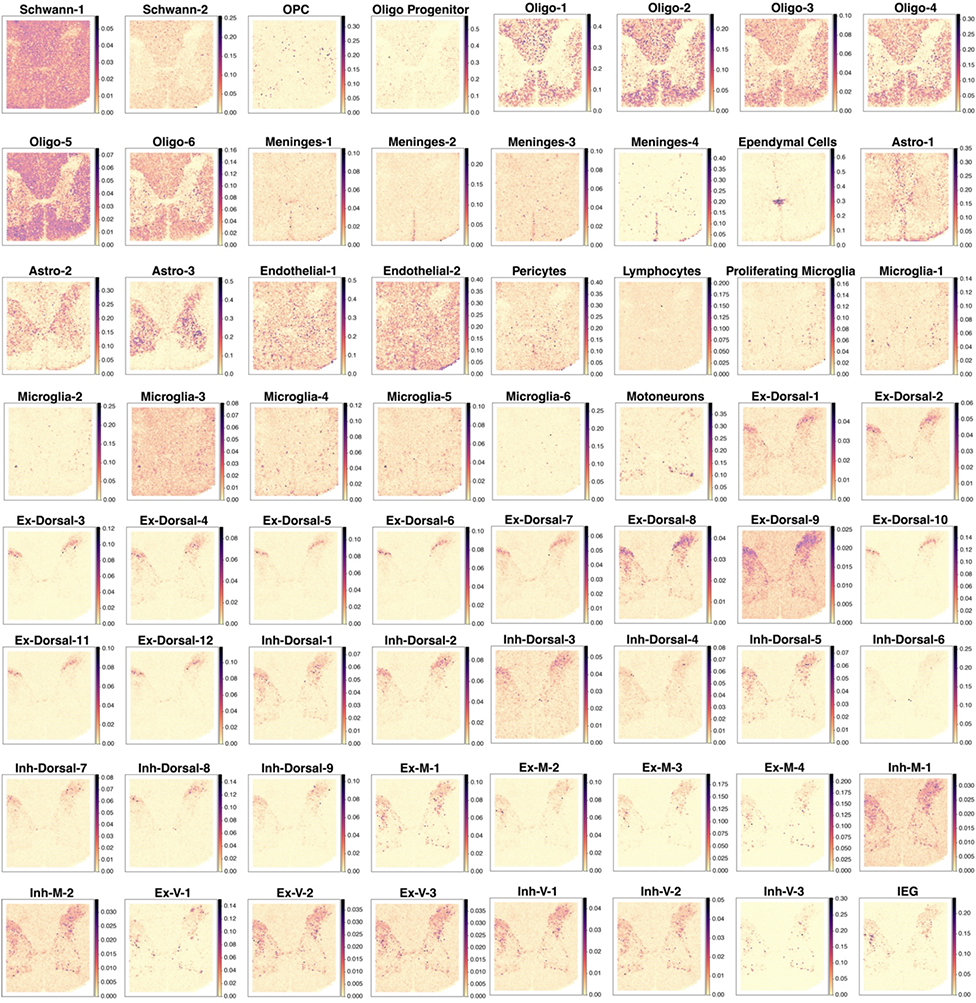
Spatial mapping of estimated cell abundances (color intensity) for 64 clusters from human spinal cord single nucleus RNA sequencing data (Cell2Location). Estimated cell abundance is colored from yellow (low) to purple (high).

**Supplemental Fig. S7.**
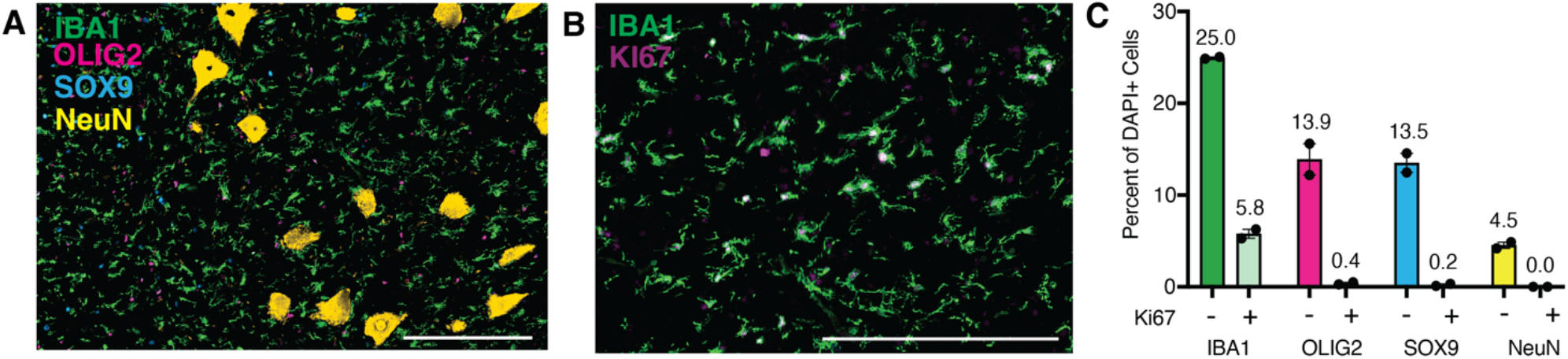
**A,** Multiplex immunohistochemistry of the lumbar human spinal cord, stained for IBA1 (green), OLIG2 (pink), SOX9 (turquoise), and NeuN (yellow). Scale bar is 250 µm. **B,** Multiplex immunohistochemistry of the lumbar human spinal cord, stained for IBA1 (green) and KI67 (purple). Scale bar is 250 µm. **C,** Bar plot showing the percent of DAPI-expressing cells in the human spinal cord that express NeuN, OLIG2, IBA1, SOX9 and KI67. Error bars are ± SEM, N=2.

**Supplemental Fig. S8.**
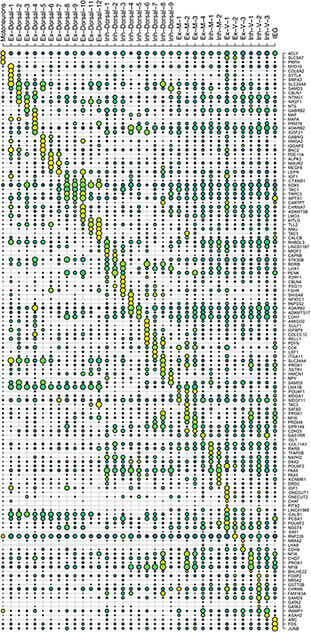
Dot plot depicting expression of genes within neuronal subpopulations. Expression is indicated by color, purple (low) to yellow (high).

**Supplemental Fig. S9.**
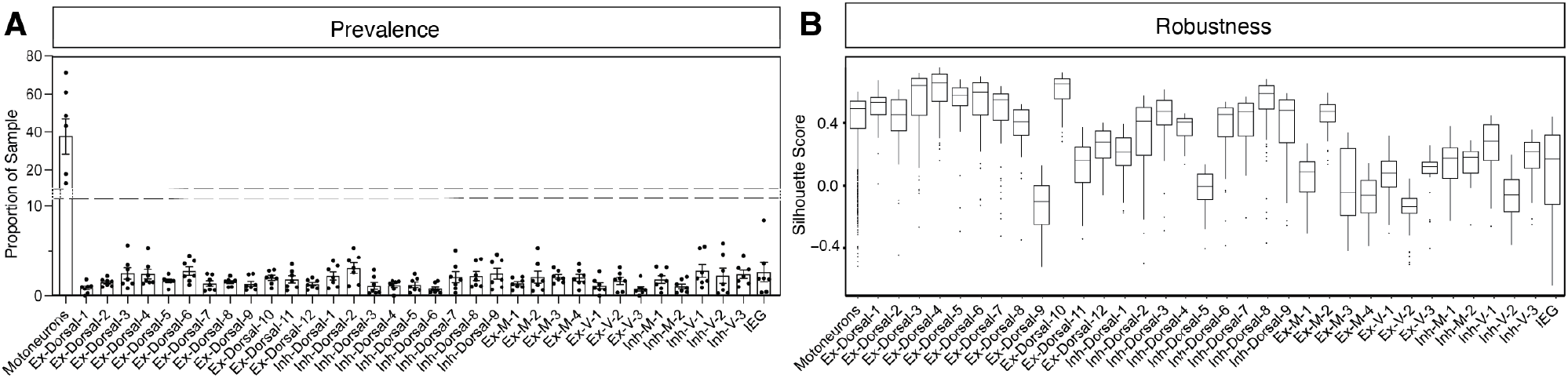
Integration and sub clustering of Neuronal sub-types in human spinal cord. **A,** Bar plot showing the proportion of a given neuronal cluster in each donor. Error bars are ± SEM, N=7. **B,** Box plot showing distribution of silhouette scores per nucleus per neuron population in order to assess cluster robustness. A high silhouette score indicates distinctiveness of a cluster.

**Supplemental Fig. S10.**
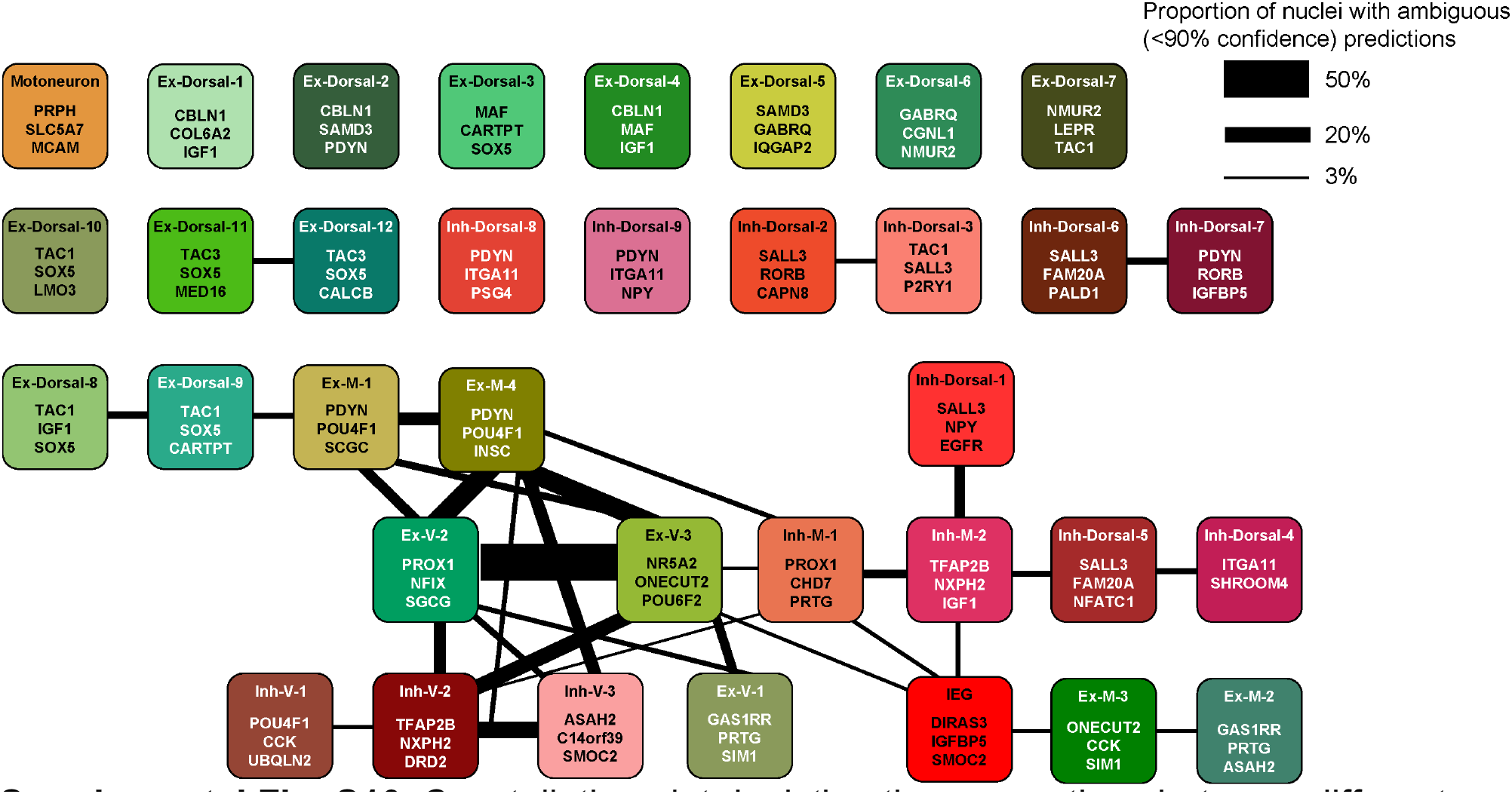
Constellation plot depicting the connections between different neuronal clusters, based on 100 iterations of post-hoc classification. The nodes are different clusters and the edges correspond to proportion of nuclei that were ambiguously predicted and are shared between the clusters during multiple iterations. The genes in each box represent a unique combination of markers to identify each cluster. Ex- Excitatory, Inh- Inhibitory, M- Mid, V- Ventral.

**Supplemental Fig. S11.**
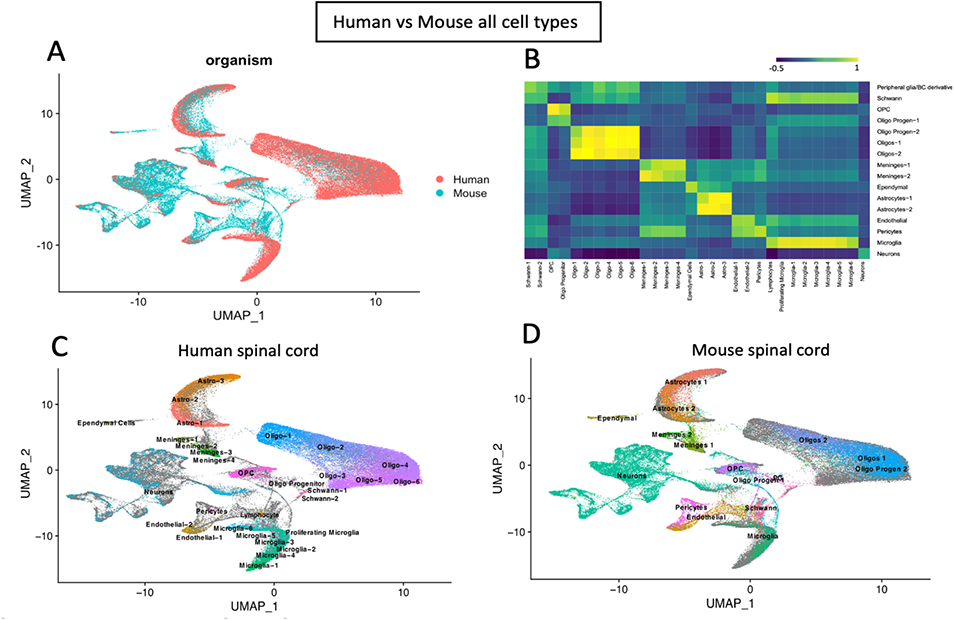
Cross species analysis between Human spinal cord and harmonized mouse spinal cord atlas. (Russ et al., 2021). **A,** Integration of the human and mouse spinal cord datasets (includes all cell types) **B,** Plot showing correlation between mouse and human cell types. Correlation is colored from purple (low) to yellow (high) and was calculated using principal components. **C,** All the human spinal cord cell types colored and labeled on integrated cross-species UMAP. Cells of the oligodendrocyte lineage are shown in blue/purple and include oligodendrocyte precursor cells (OPC), progenitors (Oligo Progen), six groups of oligodendrocytes (Oligo-1 through Oligo-6), as well as two populations of Schwann cells (Schwann-1 and –2). Microglia cells are shown in seafoam and includes a putatively proliferating population (Prolif Micro) and six groups of microglia (Micro-1 through Micro-6). Astrocytes are shown in salmon and orange and include three populations (Astro-1 through Astro-3). Meninges are shown in green and include four populations (Men-1 through Men-4). Vascular cells include two groups of endothelial cells (Endo-1 and –2) in olive and pericytes (Peri) are shown in pink. Ependymal cells are shown in khaki. Neurons are shown in teal. **D,** All the mouse spinal cord cell types colored and labeled on integrated cross-species UMAP. Cells of the oligodendrocyte lineage are shown in blue/purple and include oligodendrocyte precursor cells (OPC), two groups of progenitors (Oligo Progen-1 and Oligo-Progen-2), two groups of oligodendrocytes (Oligo-1 and Oligo-2), as well as a population of Schwann cells (Schwann-1 and –2). Microglia cells are shown in green. Astrocytes are shown in salmon and orange and include two populations (Astro-1 and Astro-2). Meninges are shown in green and include two populations (Meninges-1 and Meninges-2). Vascular cells include two groups of endothelial cells in olive and pericytes (Peri) are shown in pink. Ependymal cells are shown in khaki. Neurons are shown in teal.

**Supplemental Fig. S12.**
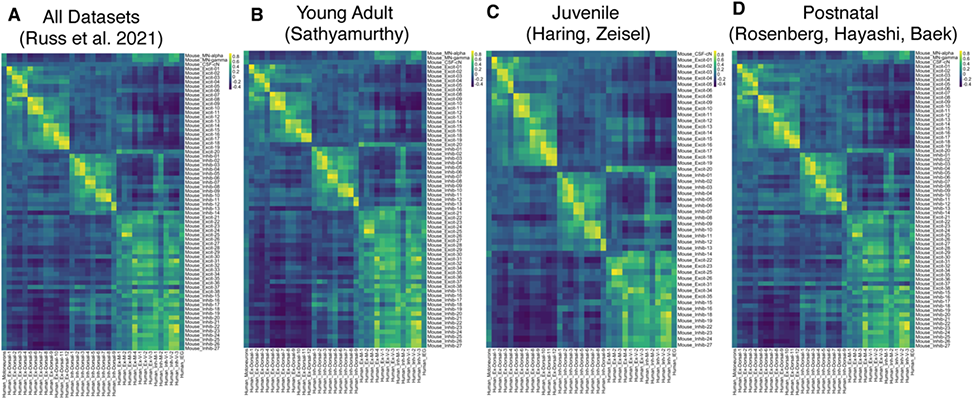
Cross species analysis of human spinal cord and harmonized mouse spinal cord neuronal subtypes based on PC 1-22. **A,** Heatmap of the human neurons vs mouse spinal cord neuronal subtypes from Russ et al. 2021. **B,** Heatmap of the human neurons vs mouse spinal cord neuronal subtypes from young adult (Sathyamurthy at el. dataset from Russ et al. 2021). **C,** Heatmap of the human neurons vs mouse spinal cord neuronal subtypes from a juvnile age (Haring and Zeisel datasets from Russ et al. 2021). **D,** Heatmap of the human neurons vs mouse spinal cord neuronal subtypes from a postnatal age (Rosenberg, Hayashi and Baek datasets from Russ et al. 2021). Correlation was calculated by principal component and is indicated by color, ranging from purple (low) to yellow (high).

**Supplemental Fig. S13.**
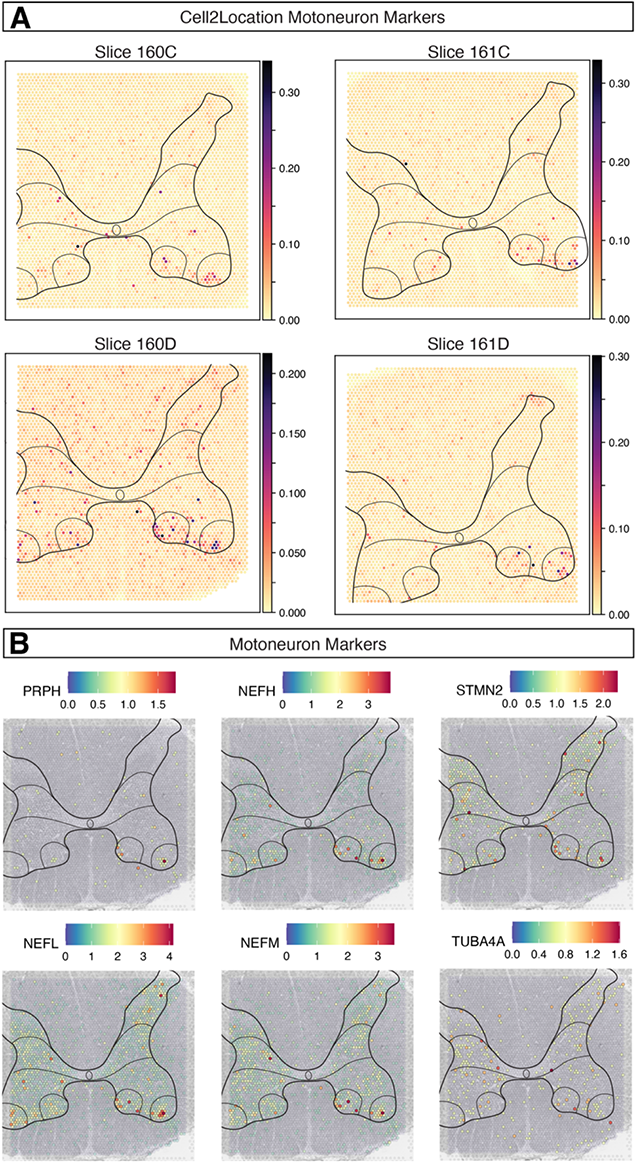
Spatial expression of human motor neuron markers. **A,** Spatial expression of Cell2Location motoneuron predicted gene signature in 4 lumbar cord sections from a single donor spinal cord. Expression is indicated by color ranging from yellow (low) to purple (high). **B,** Spatial expression of motoneuron markers in a section of lumbar human spinal cord. Expression is indicated by color, ranging from purple (low) to red (high).

**Supplemental Fig. S14.**
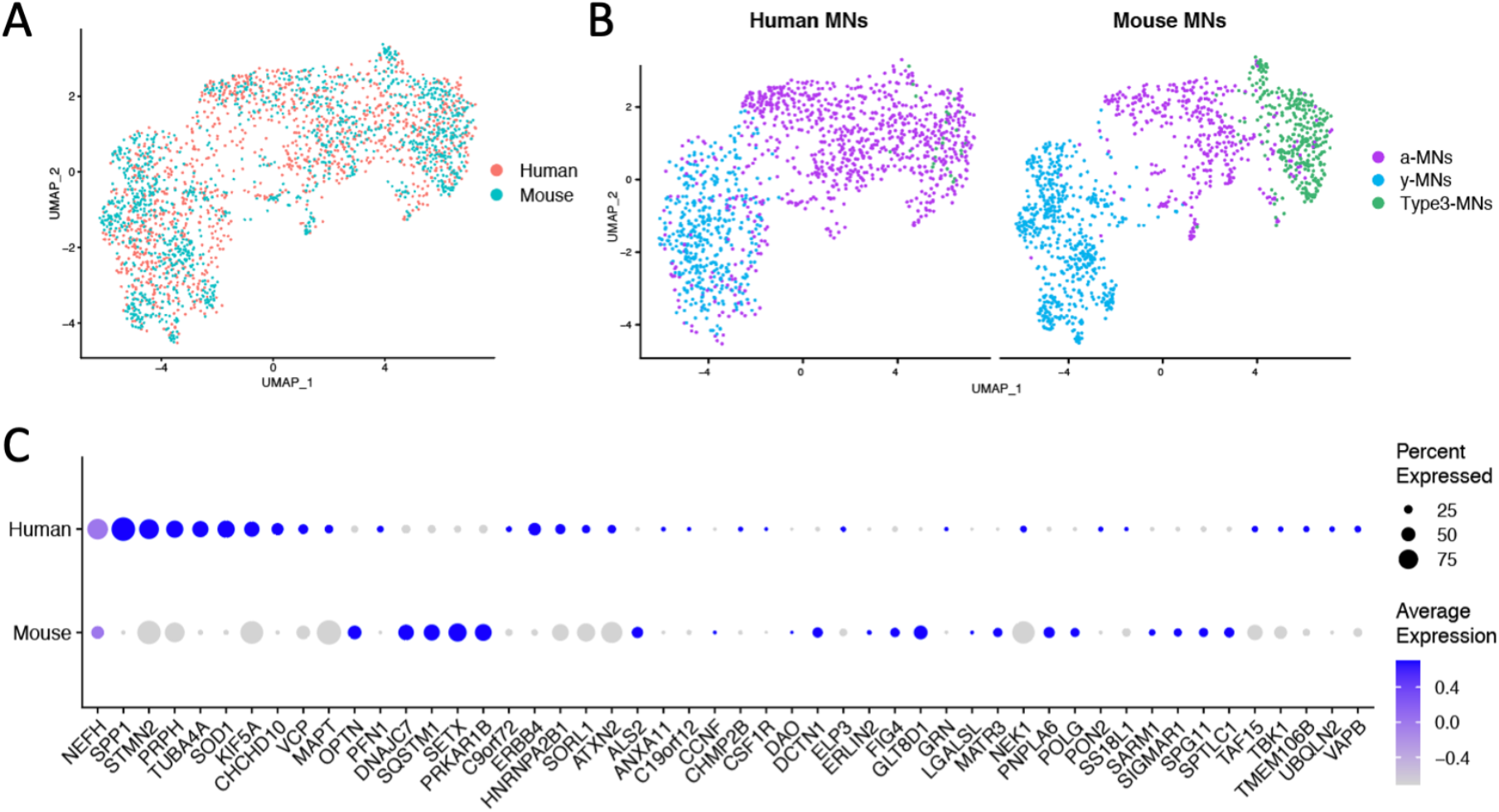
Human and mouse motor neurons differentially express ALS risk genes. **A,** UMAP representation of integrated human and mouse MN data (*46*) by dataset. **B,** UMAP representations of co-clustering of human and mouse MN data revealing potential alpha and gamma MN subtypes in human. **C,** Dot plot showing expression of known ALS risk genes across human and mouse MNs. The size of the dot corresponds to the percentage of cells that belong to particular category. The color corresponds to Average expression across all cells for a particular class.

**Supplemental Fig. S15.**
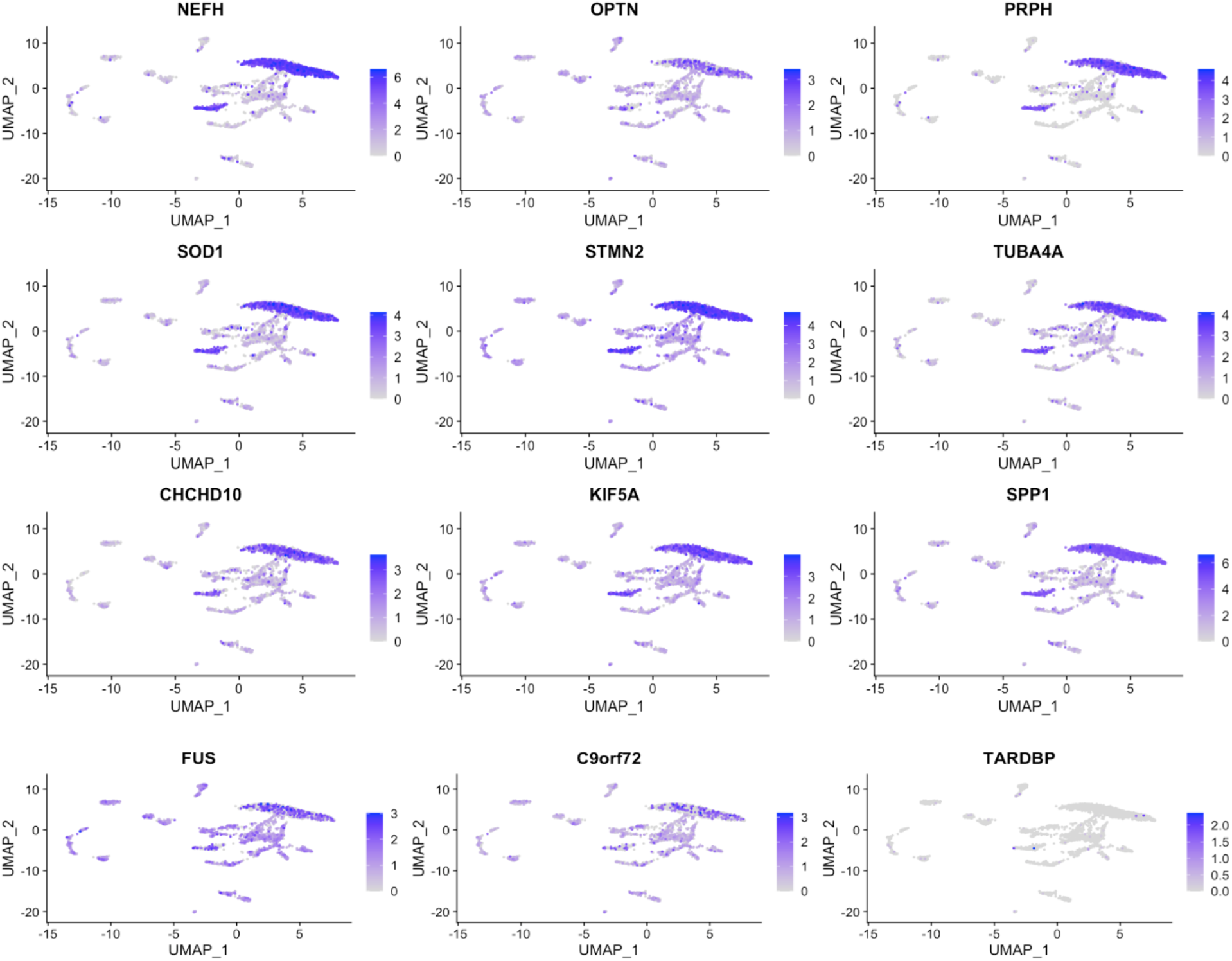
Expression of ALS-related genes in human spinal cord neurons. Log-normalized expression of NEFH, OPTN, PRPH, SOD1, STMN2, TUBA4A, CHCHD10, KIF5A, SPP1, FUS, C9orf72, TARDBP in the neurons represented in a UMAP plot. Color intensity from grey to dark blue corresponds to the amount of log normalized expression with dark blue being highest and grey being the lowest expression.

**Supplemental Fig. S16.**
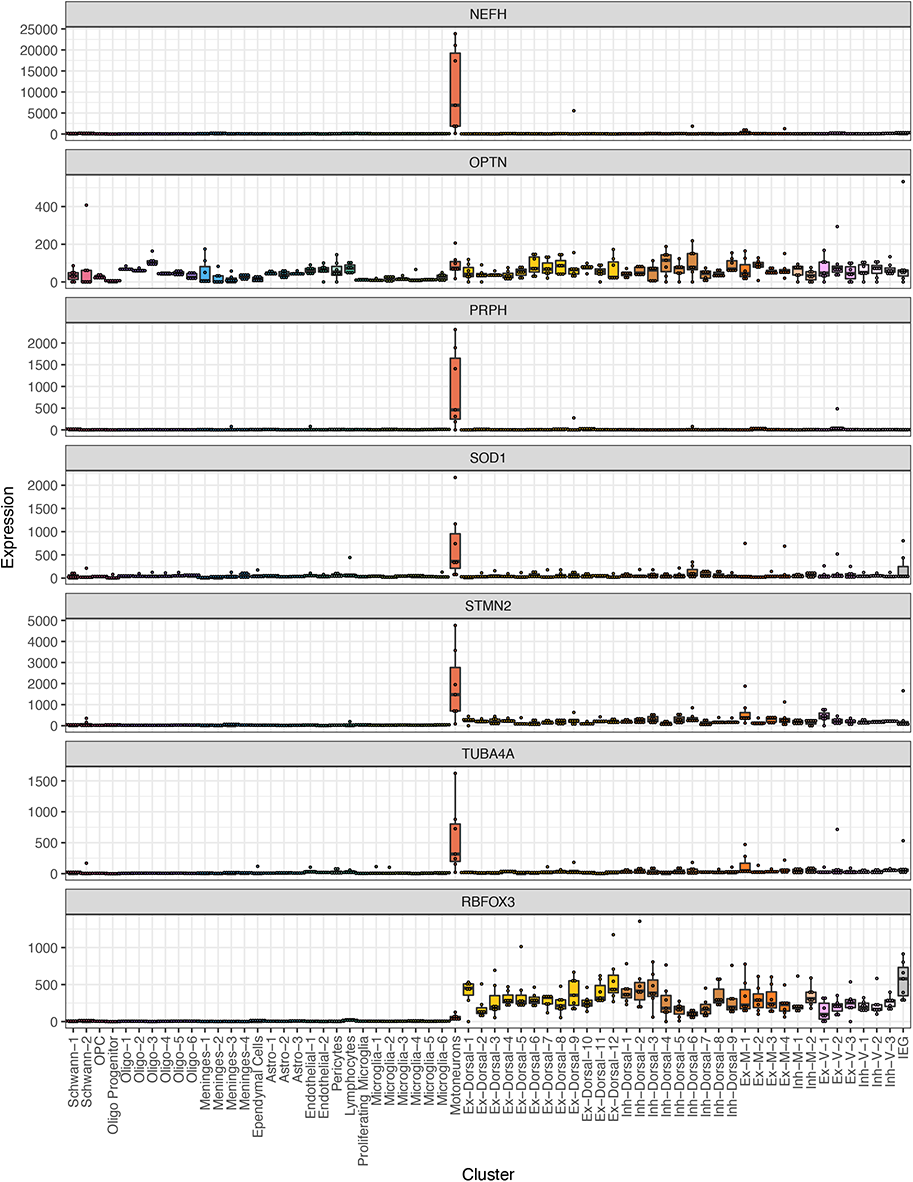
Expression of ALS-related genes in all human spinal cord cell types. Box plot shows per-cluster and per-sample expression (Counts per Million) of ALS-related genes (NEFH, OPTN, PRPH, SOD1, STMN2, TUBA4A, RBFOX3) in order to examine consistency/variability across subjects.

**Supplemental Fig. S17.**
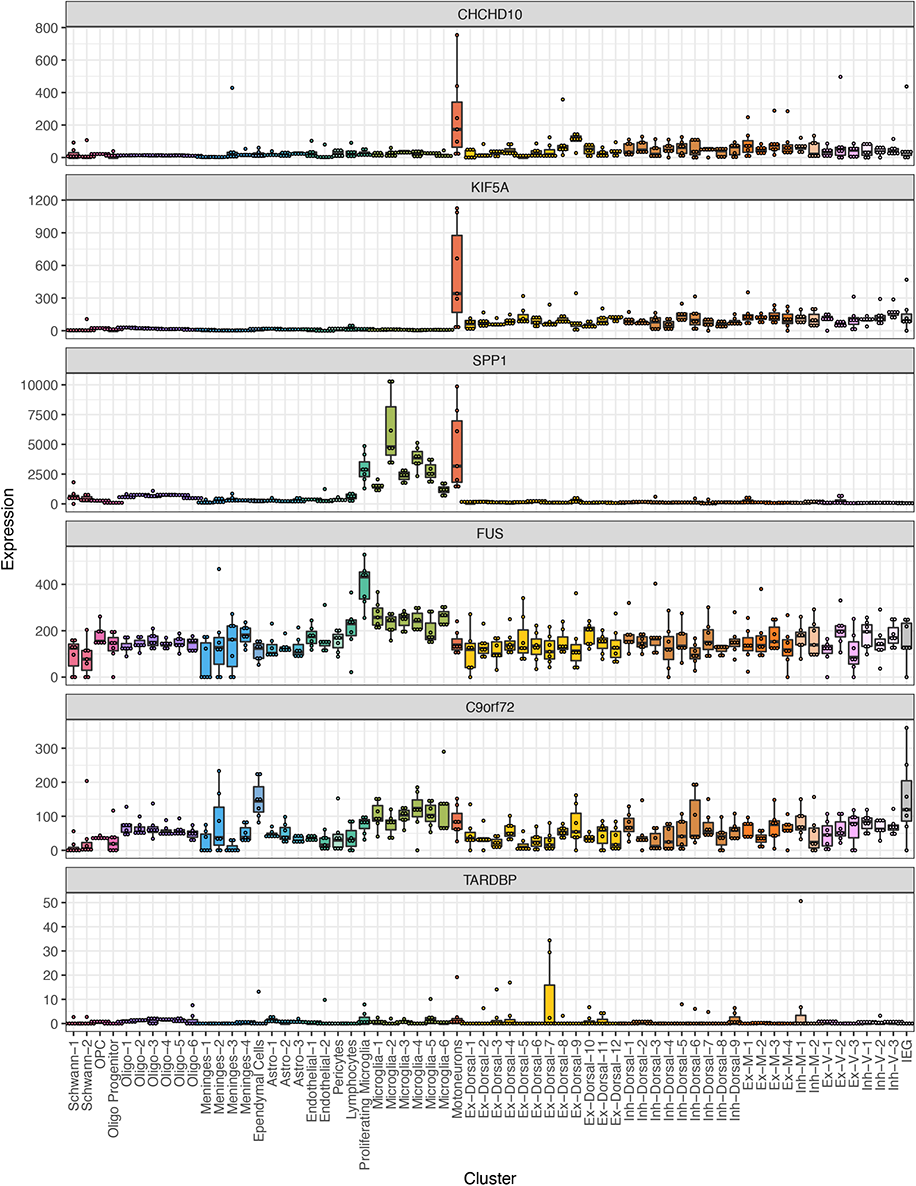
Expression of ALS-related genes in all human spinal cord cell types. Box plot shows per-cluster and per-sample expression (Counts per Million) of ALS-related genes (CHCHD10, KIF5A, SPP1, FUS, C9orf72, TARDBP), in order to examine consistency/variability across subjects.

**Supplemental Fig. S18.**
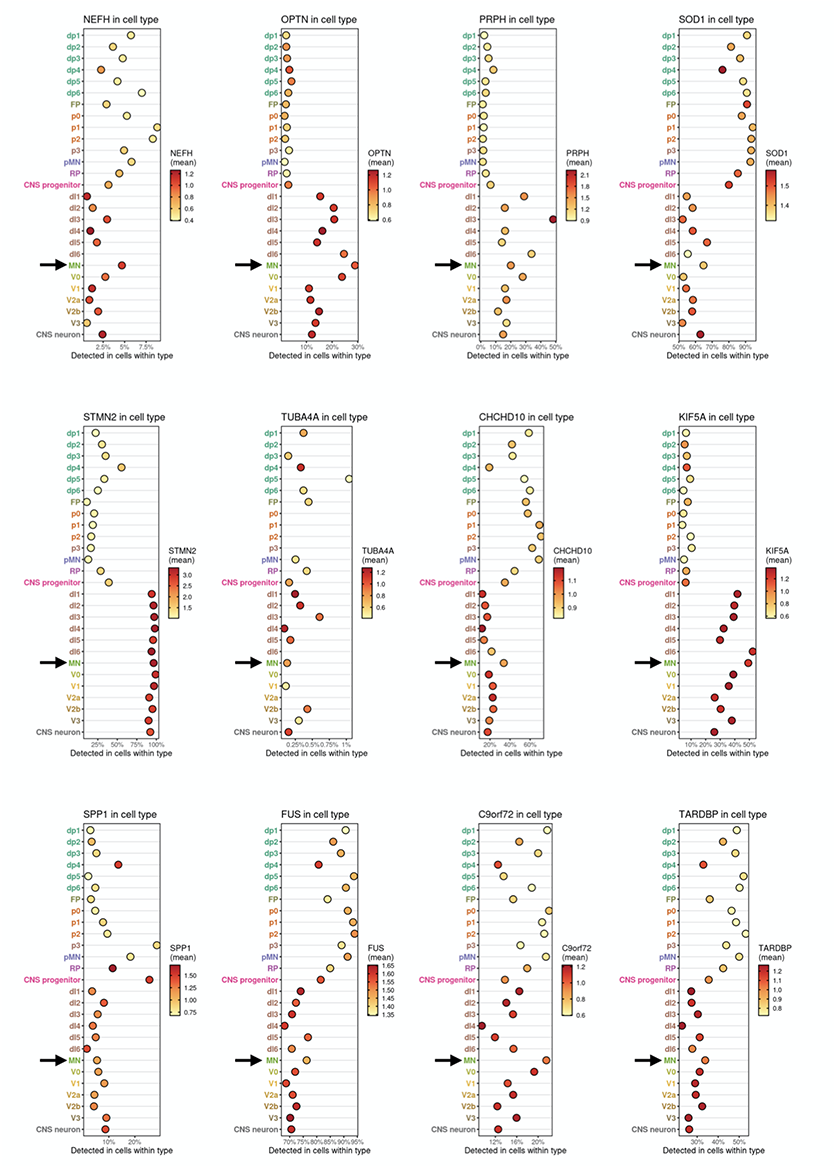
Expression of ALS-related genes in embryonic human spinal cord. Plot showing the level (color) and percent expression (location on x-axis) of 12 selected ALS-related genes in human embryonic progenitor and post-mitotic cell-types, based on https://shiny.crick.ac.uk/scviewer/neuraltube/ from (*25*). Motoneurons (MN) are indicated by a black arrow.

**Supplemental Fig. S19.**
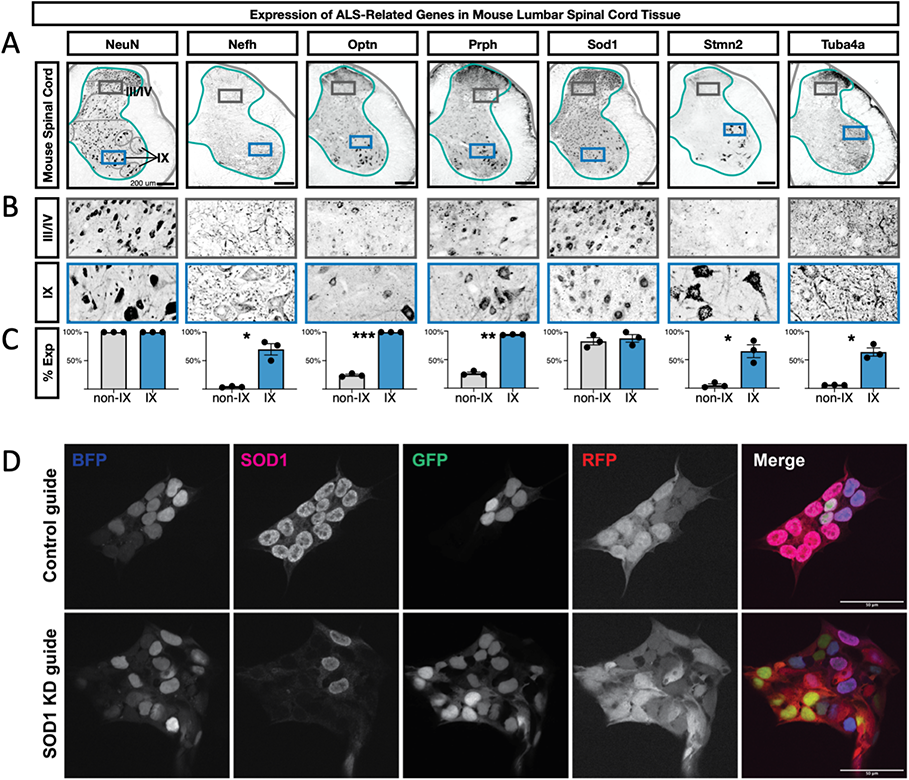
Expression of ALS-related genes in mouse lumbar spinal cord tissue. **A,** Antibody staining on lumbar spinal cord from aged mice (11 months old) for the orthologous proteins to those shown in Fig 3C in main manuscript. Gray matter outlines are shown in teal and boundaries of lamina I/II, III/IV, V/VI, VII/VIII, IX, and X are shown in gray. The boxes indicate the enlarged images in panel B. **B,** Inset of the images in panel A from the boxed region in laminae III/IV or lamina IX. Scale bars are 200 µm and the width of the enlarged images is 200 µm. **C,** Quantification of the percent of NeuN+ neurons that co-expressed the indicated proteins in either all neurons not in lamina IX (non-IX) or those in lamina IX. The mean +/-s.e.m.are shown. The plotted values and number of cells counted in each subject and category are available in Supplemental Table 5. Paired t-test results are shown where * indicates p < 0.05, ** indicates p < 0.005. **D,** Representative images of human inducible pluripotent stem cells (hiPSCs) with SOD1 knockdown and control guides 2 days after knockdown. Cells have nuclear-localizing GFP (green) and cytosolic RFP (red). BFP (blue) signifies guide uptake. Cells were stained for SOD1 (magenta). Scales bar are 50 μm.

**Supplemental Fig. S20.**
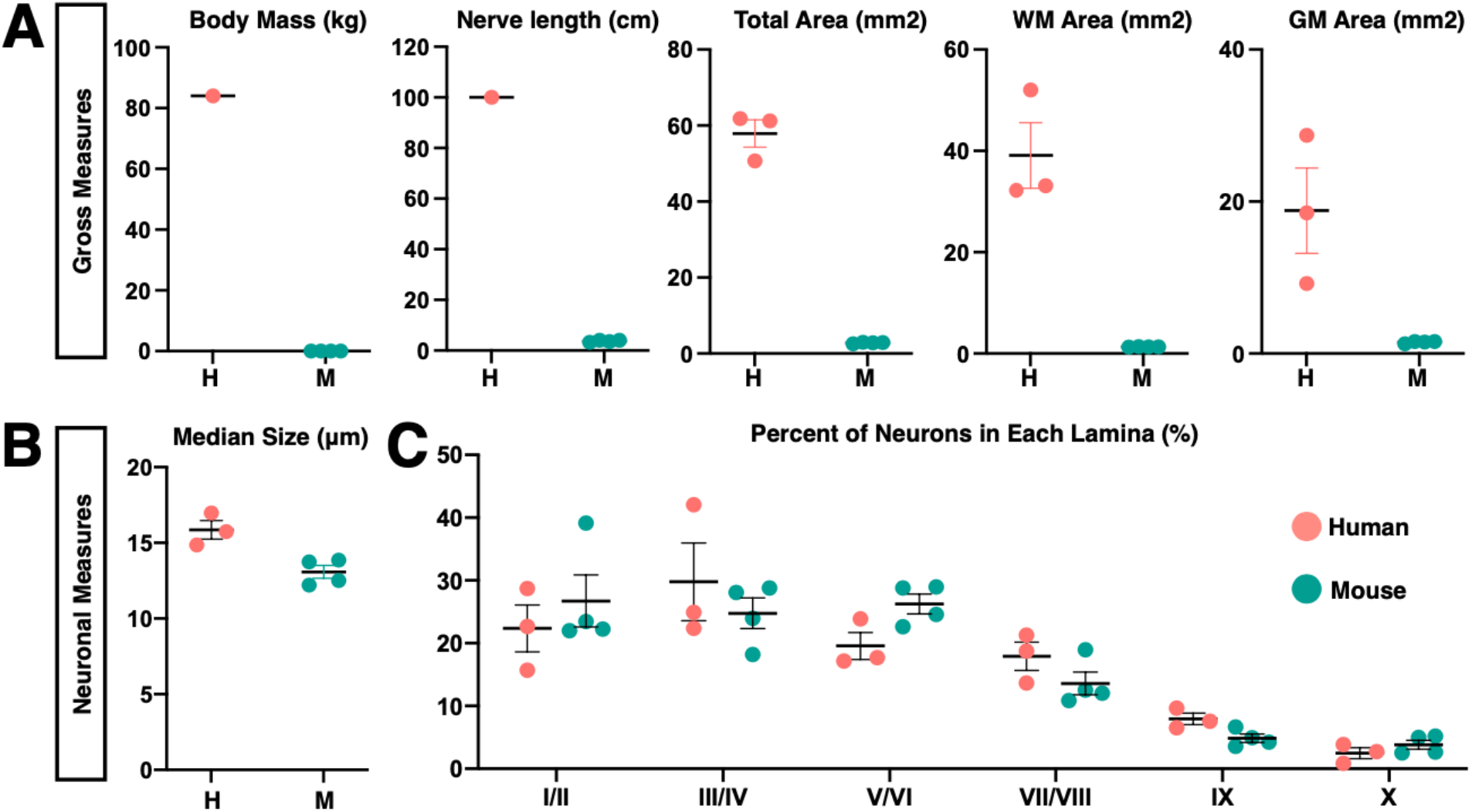
Gross anatomical and neuronal measurements of the human and mouse lumbar spinal cords. **A,** Measures of body mass, nerve length, total area, white matter (wm) area and grey matter (gm) area in the human (pink)and mouse (teal) lumbar spinal cord. Sources for human body mass (https://www.cdc.gov/nchs/fastats/body-measurements.htm) and for human nerve length (*56*). **B,** Median size of human and mouse neurons (μm). **C,** Percent of lumbar spinal cord neurons that reside in a given Rexed lamina. Error bars are ± s.e.m.

**Supplemental Fig. S21.**
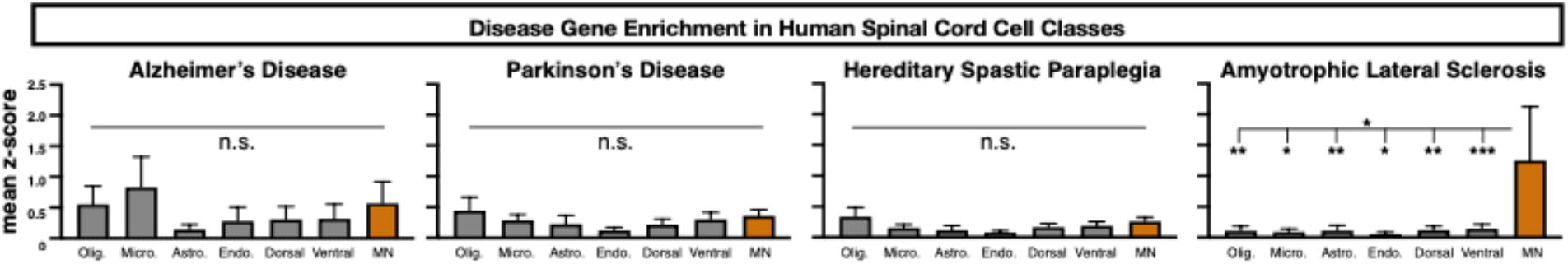
Comparison of the z-scores (mean ± s.e.m.) for genes associated with the degenerative diseases Alzheimer’s disease, Parkinson’s disease, HSP, and ALS in seven different broad classes of cells: oligodendrocytes (Olig.), microglia (Micro.), astrocytes (Astro.), endothelial cells (Endo.), dorsal horn neurons (Dorsal), ventral horn neurons (Ventral), and motoneurons (MN, orange). Gene lists for each disease are available in Data File Table S6. One-way non-parametric Friedman’s test was used to determine whether any cell types varied within each panel of disease genes and subsequently, non-parametric Wilcoxon tests were used to test each pair of cell types for significant differences. Friedman’s test p = 0.0278. * indicates p < 0.05, ** indicates p < 0.005. Error bars are ± s.e.m.

## Footnotes

1 With our current approach, the human motoneuron cluster could not be divided into more refined types. This may reflect technical limits (these nuclei contained a relatively low number of genes per nucleus) or biological continua amongst motoneuron features. Co-clustering with mouse MNs from previously published datasets suggested a division into alpha/beta and gamma sub-types but these were not clearly separated by human marker genes. As a result, human motoneurons were analyzed as one group.

